# Aducanumab Binding to Aβ1-42 Fibrils Alters Dynamics of the N-Terminal Tail While Preserving the Fibril Core

**DOI:** 10.1101/2025.11.01.686027

**Authors:** Ravi Shankar Palani, Christopher G. Williams, Dev Thacker, Robert Silvers, Fang Qian, Paul H. Weinreb, Leonard J. Mueller, Sara Linse, Robert G. Griffin

## Abstract

Aducanumab, a human IgG1 antibody with plaque-clearing effects and modest clinical benefit, binds selectively to aggregated Aβ via the N-terminal region. Yet, the molecular details of how the antibody engages Aβ_1-42_ fibrils remain unresolved. Using magic-angle spinning nuclear magnetic resonance, we show that binding of aducanumab preserves the overall architecture of the Aβ_1-42_ fibril core while inducing significant structural and dynamic perturbations in the N-terminal region. Antibody binding markedly reduces flexibility in this domain, with the appearance of sidechain resonances from residues D1, E3, and histidine (likely H6) in dipolar-based experiments. These sidechains—previously observed only in scalar-coupling spectra of the unbound state—indicate rigidification of residues that were dynamic. The interaction extends to S8 and Y10, indicating broader fibril engagement than the minimal epitope (residues 3–7) defined in fragment-based studies. Perturbations in the C-terminal segment (G37–A42) are consistent with its spatial proximity to the antibody-bound N-termini of neighboring monomers. Cryo-TEM images reveal fibrils bundling in the presence of aducanumab, consistent with lateral association via antibody cross-linking, supporting a model where surface coating and steric hindrance suppress secondary nucleation. This mode of action restricts monomer access to catalytic sites on fibril surface, resulting in partial inhibition (∼three-fold reduction) of secondary nucleation. The effect depends on high avidity and relatively high stoichiometry, but is ultimately limited by antibody size relative to N-terminal spacing along the fibril. These findings provide atomic-level insights into aducanumab’s binding mode and supply a structural framework for understanding antibody-mediated fibril recognition and for guiding next-generation therapies targeting Aβ aggregates in Alzheimer’s disease.

**Significance Statement:** Understanding how therapeutic antibodies interact with amyloid-β (Aβ) fibrils is crucial for developing effective Alzheimer’s disease treatments. Magic angle spinning NMR provides atomic-level insights into the binding of aducanumab to mature Aβ_1-42_ fibrils. Aducanumab binding preserves the fibril’s core structure but slows the dynamics of the N-terminal domain of Aβ. This interaction, which spans D1 to S8 and extends to Y10 on the fibril surface, is consistent with a mechanism in which N-terminal binding by the antibody interferes with aggregation steps like secondary nucleation. These findings detail how aducanumab engages its target fibril and provides insights relevant to other clinically approved antibodies and next-generation therapies that recognize the Aβ N-terminal region.

## Introduction

Alzheimer’s disease (AD) is the leading cause of dementia worldwide, affecting over 50 million people and posing a major social and economic burden. One of the pathological hallmarks of AD is the accumulation of extracellular amyloid plaques, predominantly composed of amyloid-β (Aβ) peptides[1, 2]. The Aβ peptides are generated from the amyloid precursor protein (APP) through sequential cleavage by β-and γ-secretases[3], yielding predominantly two isoforms: Aβ_1-40_ and Aβ_1-42_[4, 5]. Although Aβ_1-40_ is more abundant, Aβ_1-42_ aggregates more rapidly[6], exhibits greater neurotoxicity[7], and is the major species found in amyloid plaques[8]. Recent studies have revealed that in addition to these canonical isoforms, a diverse array of other Aβ variants—differing in N-and C-terminal sequences and/or posttranslational modifications—exist in human AD brain tissue[9–11]. The precise roles of these alternative isoforms in disease progression remain to be fully elucidated.

Genetic studies have strongly implicated Aβ in AD pathogenesis[12, 13]. Familial AD–associated mutations in APP and presenilin genes (PSEN1 and PSEN2) often lead to increased production of Aβ_1-42_[12], altering Aβ_1-42_:Aβ_1-40_ ratios[14, 15]. Several of these mutations cluster near Glu22, a region critical for aggregation and structurally exposed in many fibril polymorphs, and are associated with altered aggregation kinetics due to reduced electrostatic repulsion and formation of pathogenic assemblies[16–20]. These observations underscore the central role of Aβ assembly in disease progression.

Studies of the Aβ aggregation mechanism have established a multistep process involving primary nucleation, elongation, and secondary nucleation—where monomers aggregate on the surface of existing fibrils to generate new toxic species[21]. Above a critical fibril concentration, secondary nucleation becomes the dominant pathway for producing low molecular weight Aβ oligomers. This leads to auto-catalytic amplification of fibril mass and the generation of oligomeric species, which is now understood to be a major driver of toxicity[21, 22]. Critically, secondary nucleation of Aβ peptides occurs along the fibril surface[23, 24] at rare catalytic sites[25] that seem to be fibril core defects[26]. Antibodies and other proteins that bind to fibril surfaces can inhibit this process, thereby reducing the flux of oligomeric species and interfering with downstream aggregation and toxicity[22, 27]. In addition to their kinetic effects, many of the antibodies are thought to facilitate microglia-mediated clearance of Aβ aggregates, contributing to overall plaque removal and therapeutic efficacy[28, 29].

Given their role as catalysts for secondary nucleation, mature fibrils are important therapeutic targets despite being less directly toxic than oligomers or protofibrils[30]. This recognition has led to the development of monoclonal antibodies that selectively bind amyloid fibrils[31] and in the case of aggregated Aβ they include aducanumab[30], lecanemab[32, 33], and donanemab[34, 35]. These antibodies are targeted to prevent further toxic species formation or to promote immune-mediated clearance[29, 33, 35, 36]. Thus, each of these Aβ antibodies has distinct binding specificities for different Aβ assemblies and varies in efficacy.

Aducanumab, a human-derived IgG1 monoclonal antibody, was identified from pooled B cells collected from cognitively normal elderly individuals. Early clinical data indicated that aducanumab reduced brain amyloid load in a dose-and time-dependent manner and was associated with slower clinical decline[29]; however, later clinical studies failed to demonstrate a consistent cognitive benefit[37, 38]. Despite these setbacks, understanding how aducanumab interacts with Aβ_1-42_ fibrils remains important for elucidating mechanisms of antibody-mediated plaque targeting and for informing the design of next-generation immunotherapies. Structural analyses of fragmented peptides have shown that aducanumab recognizes a compact linear epitope encompassing residues 3–7 of Aβ[39]. A key characteristic of aducanumab is its pronounced selectivity for aggregated forms of Aβ over monomers. Kinetic studies by Arndt et al.[39] attribute this selectivity to a profile of weak intrinsic affinity for Aβ monomers (K_D_ in the 10 micromolar range), driven by fast association and dissociation rates, which contrasts with its strong avidity achieved through multivalent binding to epitope-rich Aβ aggregates. This binding profile, shared by other aggregate-preferring antibodies like lecanemab[40] and gantenerumab[41], is distinct from antibodies such as solanezumab, which exhibit higher affinity for monomeric Aβ and have shown less efficacy in targeting pathogenic aggregates[27, 39]. This divergence is thought to reflect a balance of intrinsic affinity and avidity: aggregate-selective antibodies such as aducanumab exhibit low affinity for monomeric Aβ but achieve strong avidity through multivalent binding to repetitive epitopes on fibrils, whereas solanezumab recognizes an epitope that is more exposed in monomers, leading to higher apparent affinity for Aβ in solution and reduced engagement with aggregates. Such aggregate selectivity is believed to minimize aducanumab’s binding to physiologically abundant monomeric Aβ, which may serve several important biological functions, thereby reducing potential off-target effects. Moreover, aducanumab has been hypothesized to “coat” the surface of Aβ_1-42_ fibrils, inhibiting secondary nucleation and downstream toxic oligomer formation, in addition to promoting clearance via immune-mediated mechanisms[27].

Structural studies of Aβ aggregates have advanced our understanding of amyloid pathology. The consensus structure of synthetic Aβ_1-42_ fibrils, formed under near-physiological conditions, involves the stacking of S-shaped monomers into parallel β-sheets, generating the characteristic cross-β amyloid architecture. High-resolution MAS NMR studies by several groups—most notably Colvin et al.[42, 43], Wälti et al.[44], and Xiao et al.[45]—have independently resolved atomic-resolution structures. In the structure reported by Colvin et al.[43], the monomer adopts a three-stranded S-shaped amyloid fold. Two monomers with mirror-image S-shaped conformations assemble back-to-back, forming a dimer stabilized by hydrophobic contacts between M35 of one molecule and Q15 and L17 of the adjacent molecule, generating two distinct hydrophobic cores and burying over a dozen hydrophobic residues. This dimer then stacks along the fibril axis in a parallel, in-register arrangement, forming the fibril spine. Small-angle scattering studies by Lattanzi et al. [46] further showed that such dimeric planes organize into a two-filament, tetrameric cross-section, reinforcing this arrangement as a low-energy structural minimum[47]. Thus, there are four monomers per fibril plane and the fibril is stabilized by the burial of a large number of hydrophobic sidechains in each filament and the pairing of two protofilaments. The presence or absence of the initiating methionine (M0) does not significantly affect this core structure, as confirmed by Silvers et al.[48]. Wälti et al. and Xiao et al. reported highly similar Aβ_1-42_ fibril structures using independent MAS NMR approaches, with minor differences mainly in the completeness of distance restraints and assignment strategies[44, 45]. These studies converge on the same overall S-shaped monomeric fold and inter-monomer packing motif, supporting a dominant and thermodynamically favored fibril polymorph under physiological *in vitro* conditions.

While recombinant Aβ_1-42_ fibrils formed *in vitro* have revealed a dominant, thermodynamically favored S-shaped fold under near-physiological conditions, cryo-EM studies of brain-derived filaments by Yang et al. have identified the co-existence of distinct morphs (Type I and Type II)[49]. These brain-derived fibrils, although differing in inter-protofilament packing and side-chain interactions, share key features—such as surface-exposed E22 and D23 residues, a β-bulge at F19 and F20, etc.—that are also present in *in vitro* models. This convergence supports the relevance of *in vitro* fibrils as a tractable platform for mechanistic studies and antibody engagement[50]. Despite progress in mapping antibody epitopes and resolving these fibril structures, the detailed molecular consequences of antibody binding to mature Aβ_1-42_ fibrils remain poorly understood. Previous studies have largely relied on peptide fragments, indirect epitope mapping, or kinetic assays. Direct interrogation of how antibody binding perturbs the structural and dynamic properties of full-length Aβ_1-42_ fibrils has been lacking.

Magic-angle spinning (MAS) NMR spectroscopy can be used to investigate the structural and dynamic effects of antibody binding on proteins [51], and in this study we use MAS to examine the binding of aducanumab to Aβ_1-42_ fibrils. The data reveals that aducanumab binding dramatically slows the dynamic behavior of the N-terminal region of the fibril, consistent with direct surface engagement. This interaction manifests as spectral perturbations for residues S8, Y10, and a histidine, along with the appearance of D1 and E3 sidechain resonances in dipolar-coupling-based experiments—signals that were previously observed only in scalar-coupling-based spectra of unbound Aβ_1-42_ fibrils. These results indicate that aducanumab binding impacts the structure and dynamics of the fibril surface at these sites, offering novel molecular insights into the antibody-fibril interaction landscape, particularly concerning the N-terminal domain of Aβ_1-42_. In addition, they have implications for understanding mechanisms of antibody-mediated effects and for the design of next-generation immunotherapeutic strategies against AD.

## Results and Discussion

Fibrils were prepared by adding aducanumab to monomeric Aβ_1-42_ with aducanumab at a 1:4.5 (aducanumab:Aβ_1-42_) molar ratio because the reported stoichiometry and high avidity[27] imply that most of aducanumab would become bound at this condition. This is also inferred from sedimentation analyses (Figure S2) and cryo-TEM imaging (Figure 1 and S3). The fibrils formed in the absence of antibody are ca 10 nm wide, and have two filaments twisted around each other with a short twist distance in agreement with previous reports [46, 52, 53]. The fibrils formed in the presence of antibody appear significantly more bundled and heavily decorated by small knobs. The bundles are in the range of 20–150 nm wide suggesting that the fibrils associate in a heterogeneous manner rather than with a defined number per bundle. The large number of knobs along all bundles suggest that the fibrils are decorated with antibody along their entire length.

**Figure 1.**
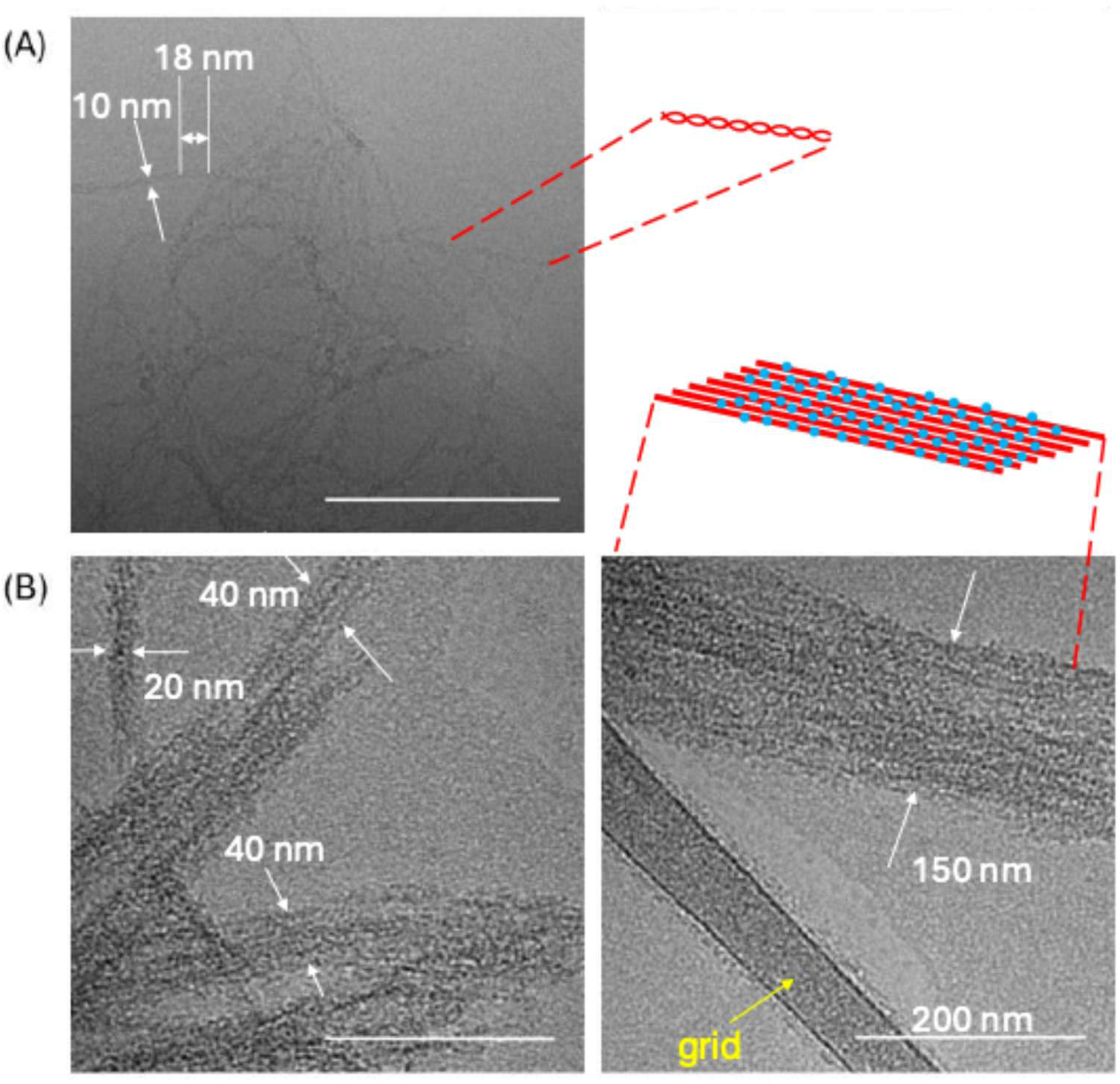
Examples of cryo-TEM images for fibrils formed from Aβ_1-42_ and Aβ_1-42_:Adu (A) One representative image of Aβ_1-42_ alone, showing a fibril width of 10 nm and period of 18 nm. The images show that the fibrils consist of two filaments twisted around one another [46, 52, 53]. This feature is illustrated with a red cartoon to the right. (B) Two images of fibrils obtained in the presence of aducanumab at 4.5:1 Aβ_1-42_:Adu molar ratio. The fibrils appear laterally assembled in bundles of 20-150 nm width and the Adu is seen as dark dots dispersed approximately uniformly along and across the bundles. This is illustrated in the cartoon with each red line representing one fibril in the bundle and the blue dots the Adu antibody. Each scale bar corresponds to 200 nm.

Fibrils were formed by adding aducanumab to freshly prepared, monomeric Aβ_1-42_ rather than adding the antibody to pre-formed fibrils as this enables us to distinguish among possible outcomes of antibody interaction: (i) fibrils forming independently with free antibody in solution, (ii) fibrils of unchanged core structure with antibody bound to the surface, or (iii) fibrils in which antibody perturbs or integrates into the core. If instead antibody were added only to mature fibrils, it would not be possible to assess outcome (iii), since the slow dissociation of Aβ_1-42_ fibrils precludes core remodeling. In the following sections, we evaluate which of these states is most consistent with the NMR data.

We compared MAS NMR spectra of Aβ_1-42_ fibrils formed in the presence of aducanumab with reference spectra from native Aβ_1-42_ fibrils formed in its absence (Ref. [48]). Figure 2 (A,C) show the ^13^C and ^15^N CPMAS spectra, respectively, of Aβ_1-42_ fibrils in the presence and absence of aducanumab. The spectra display strong overall similarity, indicating that aducanumab binding does not induce significant structural changes within the rigid fibril core comprised of residues Q15-A42. The major chemical shift features, characteristic of the S-shaped amyloid fold, remain preserved, including signals from key residues involved in the hydrophobic core and inter-monomer packing. There are, however, a few subtle differences between the spectra of fibrils formed in the presence and absence of aducanumab, most of which are localized to regions outside the well-ordered core. These minor deviations become more apparent in the multidimensional correlation spectra described later in the text. Overall, the CPMAS data confirm that the fibril core is structurally robust to antibody binding, thus excluding option (iii), enabling focused investigation of the antibody-induced changes in the dynamic and solvent-exposed regions of the fibril.

**Figure 2.**
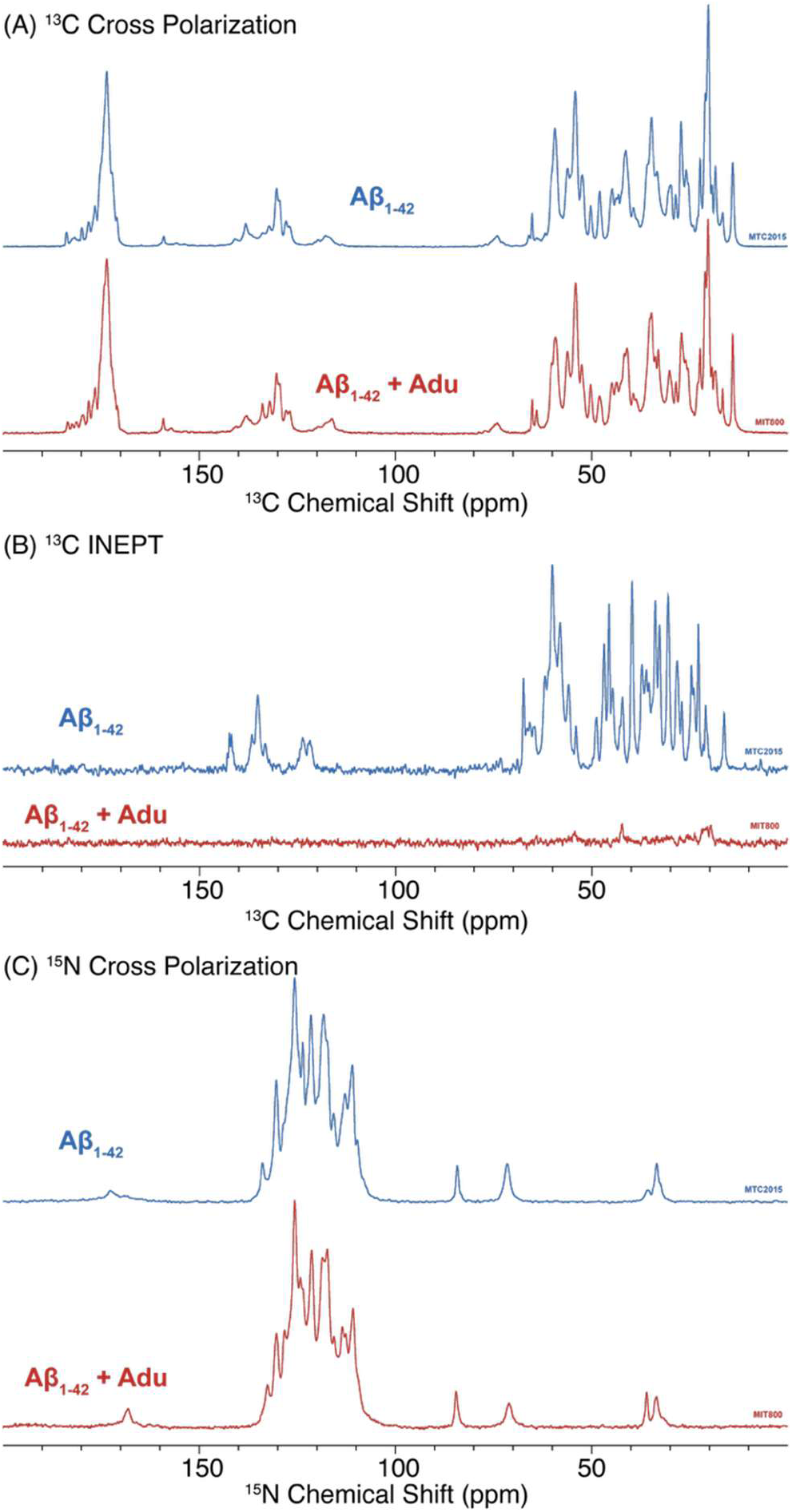
(A) ^13^C CPMAS, (B) ^13^C INEPT, and (C) ^15^N CPMAS spectra of (blue) pure, unbound U-^13^C,^15^N-Aβ_1-42_ fibrils and (red) fibrils prepared in the presence of aducanumab U-^13^C,^15^N-Aβ_1-42_ fibrils. The data were collected at 800 MHz, ω_r_/2π=20 kHz, and at 277 K. The data for the unbound Aβ_1-42_ samples are reproduced with permission from Ref.[48]. A major difference in the Aβ_1-42_ and Aβ_1-42_:Adu spectra is visible in (B) where upon binding Adu the spectrum due to the mobile residues disappears. There are other subtle differences for example in (A) the carbonyl, aromatic and serine regions of the ^13^C MAS spectra are changed by Adu binding. In the ^15^N spectra there are alterations in the His and-NH_3_^+^ regions of the spectra. These are discussed in the text.

Figure 2B shows the ^13^C MAS INEPT spectra of Aβ_1-42_ fibrils with and without aducanumab. INEPT-based detection highlights dynamic regions of the fibril that undergo fast molecular motions, typically with correlation times on the order of nanosecond or faster, enabling efficient scalar-coupling-based polarization transfer. Such regions are typically the solvent-exposed, flexible segments. In the unbound sample, the INEPT spectrum displays numerous sharp resonances, consistent with a highly mobile N-terminal domain. Upon aducanumab binding, the majority of these signals vanish, indicating that the corresponding residues undergo a substantial loss in mobility. This dramatic attenuation suggests that aducanumab binding stabilizes the N-terminal tail, either by direct interaction or by restricting local dynamics through surface association. Motions that are slowed beyond ∼10 ns may fall into a detection gap: if the dynamics are slower than tens of microseconds to milliseconds, dipolar-based recoupling methods such as CP or RFDR can detect these sites. However, residues fluctuating on the ∼100 ns to ∼10 μs timescale often experience intermediate exchange broadening and may escape detection by both scalar-and dipolar-based approaches. Thus, the attenuation of INEPT signals suggests a substantial slowing of N-terminal dynamics, possibly into this “dark” regime. As we show later, however, certain residues— specifically D1 and E3—do reappear in dipolar-based RFDR spectra, indicating that their motions are slowed sufficiently to reenter the dipolar-detectable regime. These findings support a model in which aducanumab alters the dynamic landscape of the fibril surface without perturbing the rigid core, i.e., option (ii) of the possibilities. Consistent with this interpretation, no residues retain mobility detectable by INEPT in the aducanumab-bound state (Figure S5). Even side chains that appear more intense in direct polarization than in cross-polarization are not observable in INEPT, indicating that residual epitopes do not remain freely flexible but instead fall into an intermediate motional regime that escapes INEPT detection.

The one-dimensional CPMAS and INEPT spectra shown in Figure 2 for Aβ_1-42_ fibrils in the presence of aducanumab were recorded at 800 MHz using a standard 3.2 mm MAS rotor with an ∼30 μL volume, under the same experimental conditions as the blue trace from Ref. [48] corresponding to unbound Aβ_1-42_. However, due to the molecular weights of aducanumab (142 kDa) and Aβ_1-42_ (4.5 kDa), and using the established 1:4.5 binding stoichiometry of aducanumab to Aβ_1-42_[27], the sample containing the antibody-bound fibrils has approximately 8-fold fewer moles of Aβ_1-42_ in the rotor. As a result, the overall sensitivity is substantially reduced and insufficient to support multidimensional correlation experiments, as illustrated in Figure S6. To overcome this limitation, we acquired multidimensional spectra at 600 MHz using a MAS CryoProbe, which offers a 4–5x sensitivity boost[54, 55] and a MAS rotor with a 90 μL volume, thereby enabling detection of weaker signals in the antibody-bound sample.

Figure 3 shows 2D ^13^C–^13^C 1.6 ms RFDR spectra acquired using the MAS CryoProbe. The Aβ_1-42_ fibrils in the presence (red) and absence (blue) of aducanumab show largely overlapping cross peaks in the aliphatic and carbonyl regions, indicating that the overall fibril architecture remains unchanged upon antibody binding. The excellent agreement in both peak positions and intensities indicates that the rigid core of the fibril is structurally unaffected. At the same time, the sensitivity enhancement provided by the cryoprobe enables clear detection of more subtle differences in surface-exposed residues. We observe two new cross peaks in the RFDR spectra of Aβ_1-42_ fibrils prepared in the presence of aducanumab, that are not observable in the unbound state. These peaks correspond to D1Cβ-Cγ and E3Cγ–Cδ correlations, contacts involving the sidechain carboxyl groups. Notably, the corresponding D1Cβ and E3Cγ chemical shifts closely match values reported in scalar-coupling–based INEPT-TOBSY spectra of unbound Aβ_1-42_ fibrils (Ref. [42]), where sidechains are highly mobile. In our case, the disappearance of INEPT signals combined with the appearance of these RFDR cross peaks indicates that the sidechains of D1 and E3, previously dynamic in the unbound state, become more rigid upon aducanumab binding. This observation is consistent with previous findings that p3-Aβ_1-42_ (Aβ with a pyroglutamic acid at position 3), which lacks a free E3 sidechain, binds aducanumab with significantly reduced affinity, suggesting that E3 may directly contribute to the antibody interface[39].

**Figure 3.**
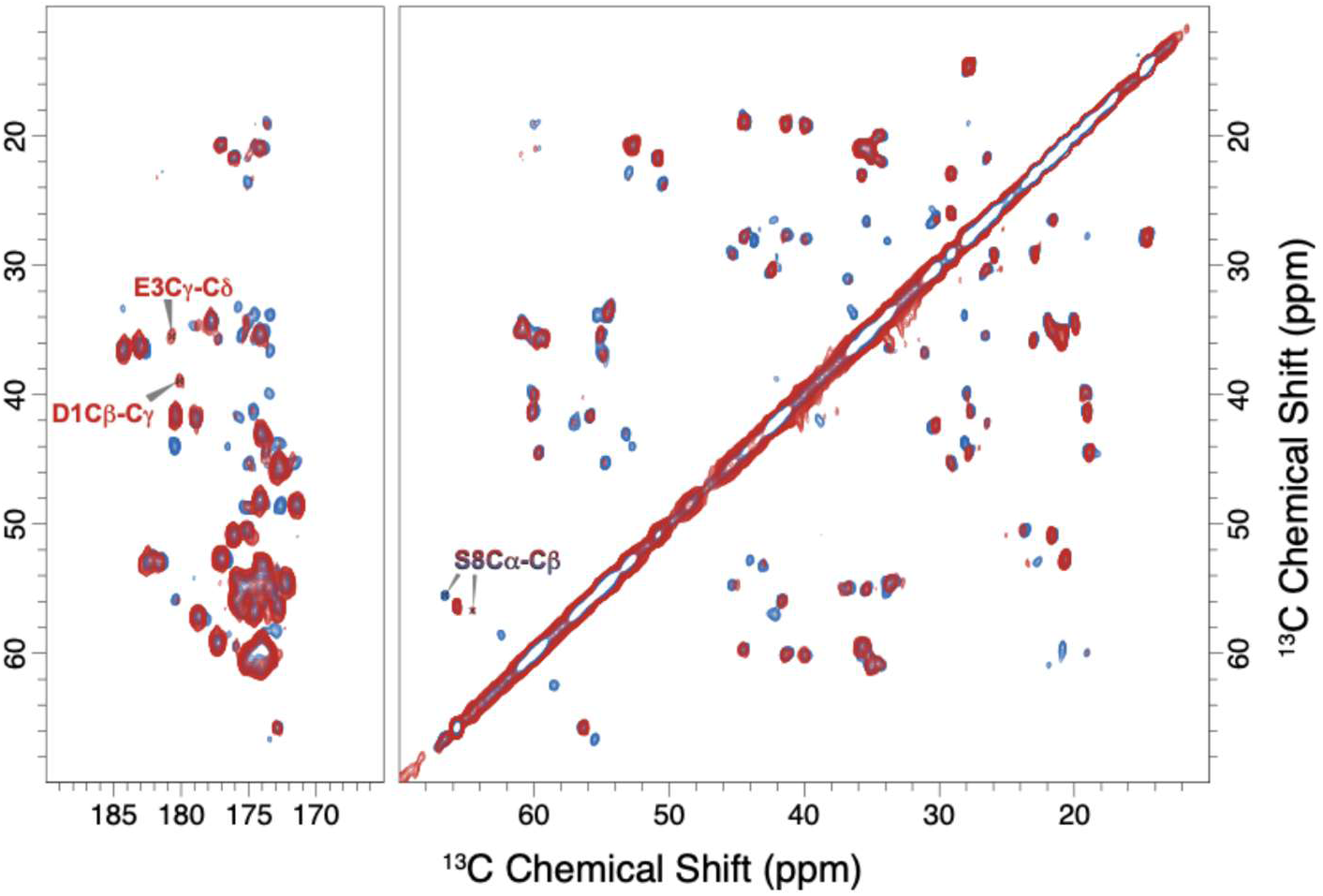
1.6 ms RFDR spectra of (red) U-^13^C,^15^N-Aβ_1-42_ fibrils formed in the presence of aducanumab recorded at 600 MHz, ω_r_/2π=13.5 kHz, and 277 K. (blue) Data for pure U-^13^C,^15^N-Aβ_1-42_ fibrils, recorded at 800 MHz, ω_r_/2π=20 kHz, and at 277 K, reproduced from Ref.[48] with permission.

Figure 4 presents 2D ^13^C–^13^C 100 ms DARR spectra highlighting differences in the fibril sample prepared in the presence of the antibody. These spectra provide insight into the local structural consequences of aducanumab binding. Notably, we observe strong perturbations of cross peaks involving residue S8 (S8Cβ–CO, S8Cα–Cβ). S8 is located immediately after the established aducanumab epitope, which encompasses residues 3-7 of Aβ ([39]). Notably, Ser8 displays a minor second set of cross peaks in unbound fibrils, suggesting a second conformation (Table S1 for assignment). These additional resonances vanish upon aducanumab binding, suggesting that the N-terminal region of the fibril participates directly in the interaction. While the main sidechain and backbone resonances for E22 (E22Cα–Cδ, E22Cγ–Cδ, E22Cβ–Cδ) appear largely unperturbed compared to reference spectra of unbound fibrils, a new cross peak involving E22Cδ is discernible in the antibody-bound sample, potentially assignable to an E22Cδ-A21Cβ interaction. The appearance of this weak peak, which is not observed in the reference spectra from Silvers et al.[48], may simply reflect the higher sensitivity of the MAS CryoProbe used in the current study. Given that A21 and E22 are adjacent residues, such a contact is structurally plausible within the fibril monomer and may not necessarily indicate a significant, antibody-induced conformational rearrangement. E22 is a known mutation hotspot in familial Alzheimer’s disease, with several variants with lower net negative charge such as Dutch (E22Q), Italian (E22K), and Osaka (E22Δ) mutations reported to significantly alter Aβ aggregation kinetics, fibril structure, and pathogenicity[16, 17, 19, 20]. This residue, located in the surface-exposed lower bend of the S-shaped fibril fold and not structurally far from the interacting N-terminal tail, appears largely unperturbed by aducanumab binding.

**Figure 4.**
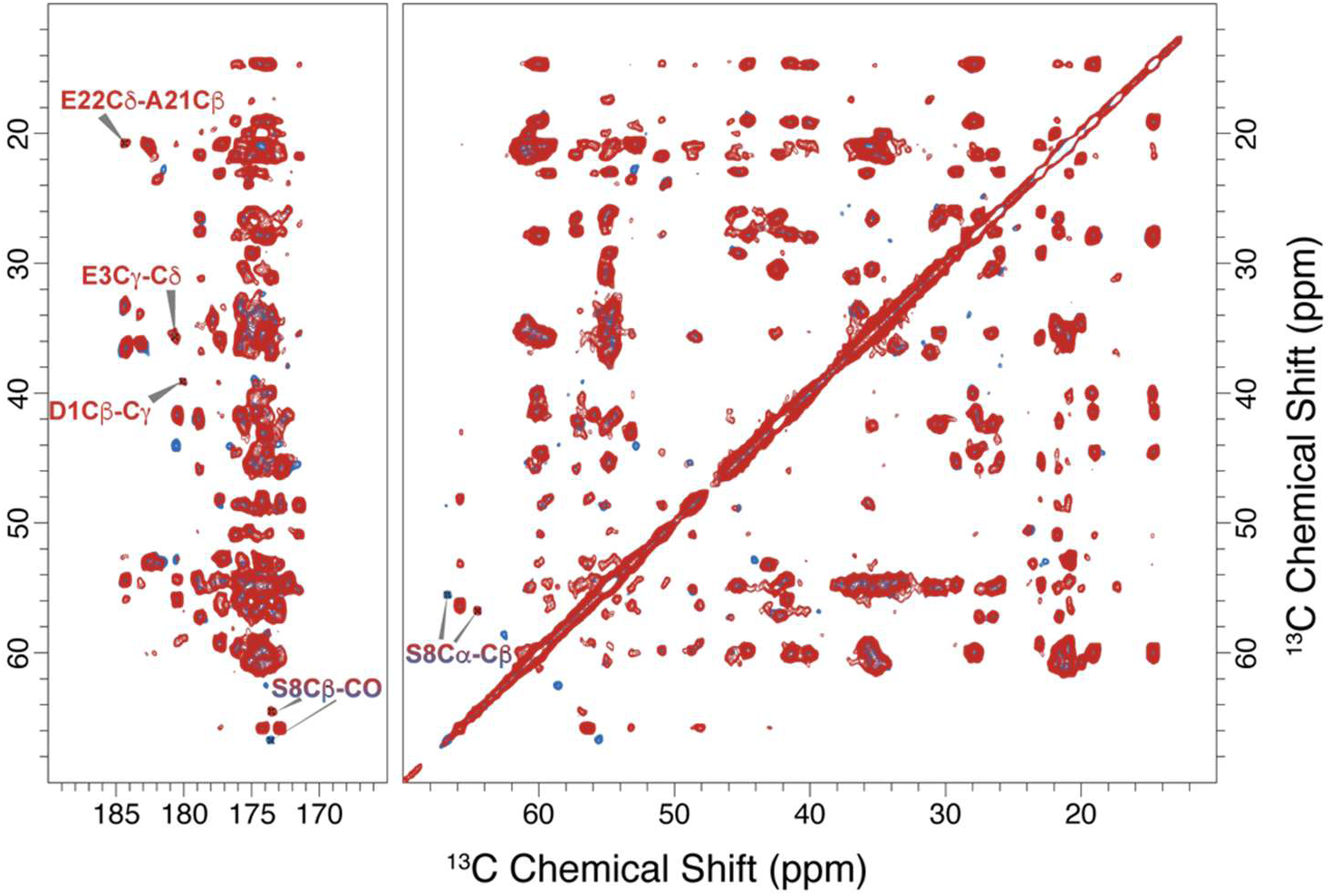
100 ms DARR spectra of U-^13^C,^15^N-Aβ_1-42_ fibrils prepared in the presence of aducanumab (red) recorded at 600 MHz, ω_r_/2π=13.5 kHz, and 277 K. Data for unbound U-^13^C,^15^N-Aβ_1-42_ fibrils (blue), recorded at 800 MHz, ω_r_/2π=20 kHz, and at 277 K, was reproduced from Ref.[48] with permission.

In Figure 5, we analyze sidechain-specific changes observed in ^13^C and ^15^N CPMAS spectra recorded using the MAS CryoProbe, which further illuminate the impact of aducanumab on the N-terminal segment. The ^13^C spectra (Figure 5A) reveal perturbations in the resonance of the Cζ carbon of Y10’s aromatic ring, along with the appearance of cross-peaks between Cζ and Cγ/Cδ in the 100 ms DARR spectrum inset, which were not observable in the unbound fibrils. These altered spectral features suggest increased rigidity or a change in Y10’s local chemical environment, indicating that the effects of aducanumab binding extend beyond the canonical epitope of residues 3-7[39]. Additional insights come from the ^15^N CPMAS data (Figure 5B), which show new or enhanced signals from the imidazole rings of His6, His13, and/or His14 in the fibril sample prepared in the presence of the antibody. While at this point our data do not allow unambiguous assignment of individual histidines, the increased definition and intensity of these sidechain resonances suggest a marked reduction in conformational flexibility for one or more histidines in the N-terminal segment with His6 being the most likely candidate. Taken together, these findings support a model in which aducanumab binding not only engages its core epitope but also restricts dynamics and modifies the local solvent-exposed surface of Aβ_1-42_ fibrils over a broader portion of the N-terminal region.

**Figure 5.**
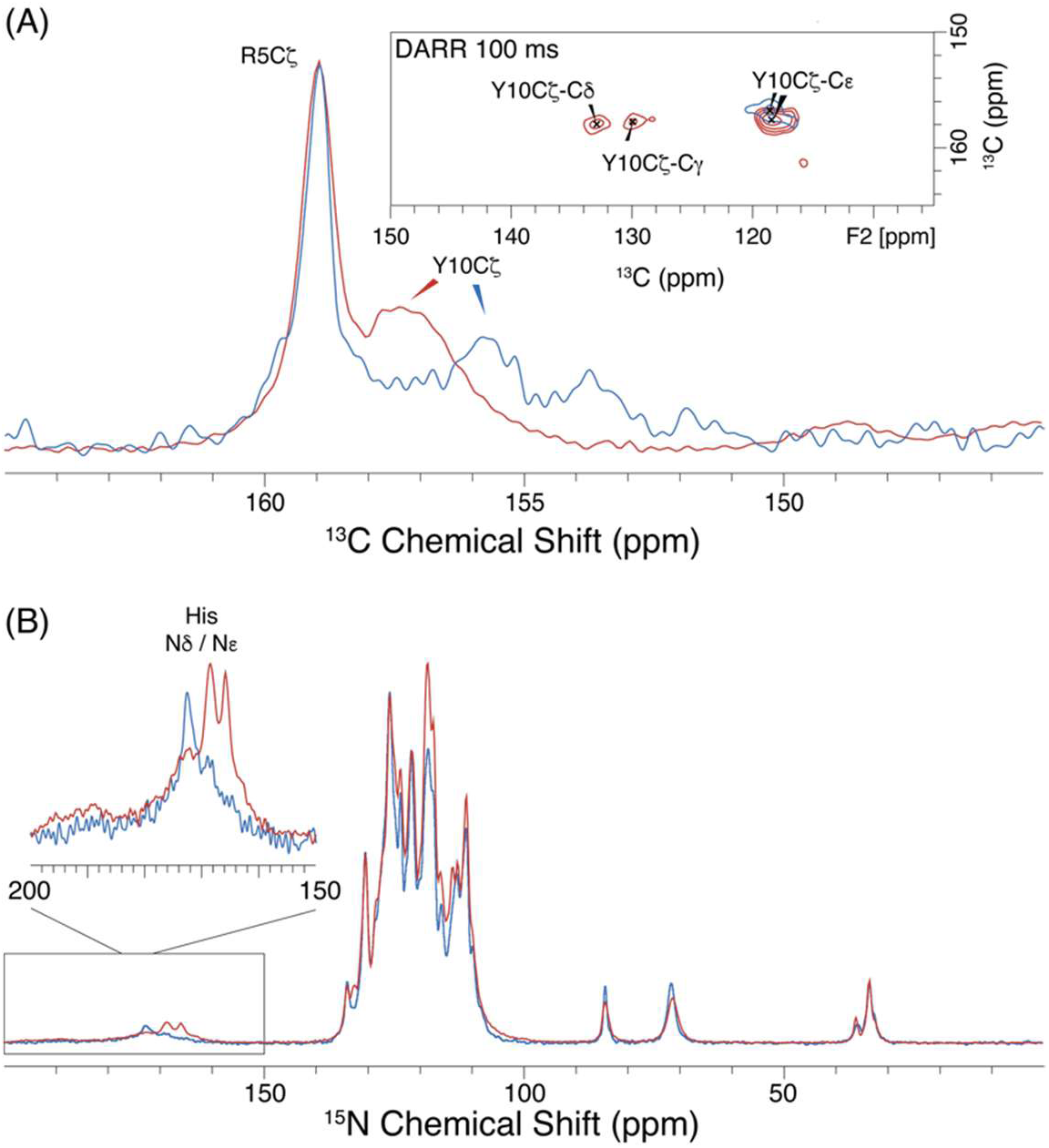
(A) ^13^C CPMAS spectra of U-^13^C,^15^N-Aβ_1-42_ fibrils prepared in the presence (red) and absence (blue) of aducanumab showing sidechain Cζ carbon resonances of R5 and Y10, while the inset shows Y10Cζ’s cross peaks with Cε, Cγ, and Cδ carbons in the 100 ms DARR spectra. (B) ^15^N CPMAS spectra of the samples with the inset showing an enlarged spectra of the His (H6, H13, H14) sidechain resonances. The data shown here of Aβ_1-42_ fibrils prepared in the presence of aducanumab (red) were recorded at 600 MHz, ω_r_/2π=13.5 kHz, and 277 K, whereas the data of unbound Aβ_1-42_ shown here is reproduced with permission from Ref.[48] and was recorded at 800 MHz, ω_r_/2π=20 kHz, and 277 K.

Comprehensive ^13^C and ^15^N chemical shift assignments for Aβ_1-42_ fibrils, both in the presence and absence of aducanumab, together with a residue-specific comparison of chemical shift differences, are provided in the Supporting Information (Tables S1–S3). The perturbations are additionally plotted in Figure 6D highlighting that the N-terminal region is perturbed while the core remains unchanged. Minor perturbations are also observed for residues G37-A42 in the C-terminal region. Since this C-terminal segment faces outward and lies in close spatial proximity to the N-terminal region of the other monomer in the filament dimer, such effects are structurally feasible and consistent with aducanumab engaging these neighboring surfaces.

**Figure 6.**
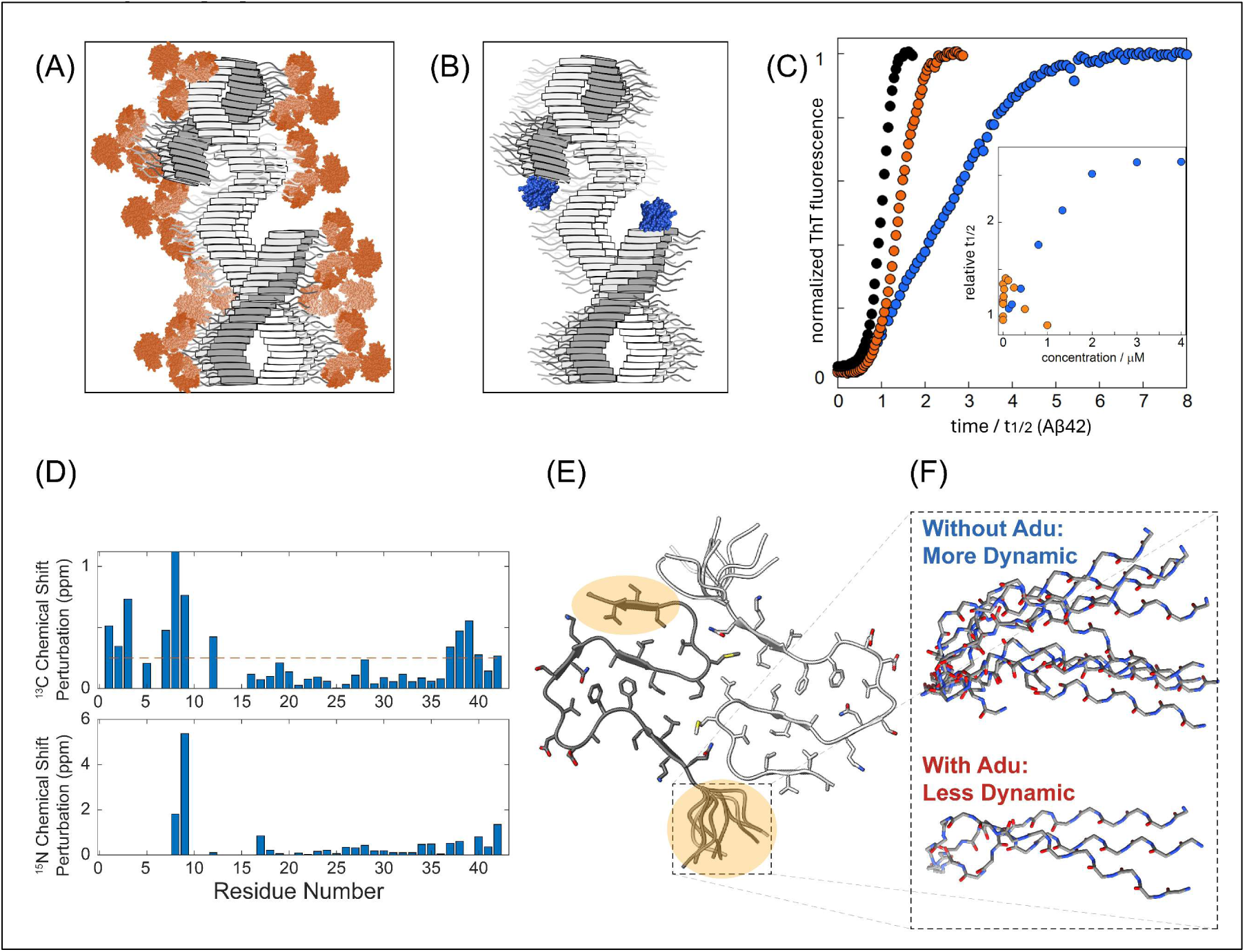
Cartoon illustrating (A) the binding of aducanumab (orange) to two N-termini per antibody with high avidity and coating of the fibril surface, hindering access to catalytic sites at fibril defects (here illustrated as dislocations, although the precise nature of defects remains to be determined), (B) the binding of Brichos (blue) to fibril defects, and (C) Aggregation kinetics data for 3 µM pure Aβ_1-42_ (black) and in the presence of 125 nM aducanumab (orange) or 2 µM Brichos (blue), shown as normalized ThT fluorescence versus time/t1/2 of pure Aβ_1-42_, replotted from refs. [22, 27]. The inset in panel C shows the effect on t_1/2_, the time at which half the monomers have converted to fibrils for aducanumab (orange, Ref. [27]) and Brichos (blue, Ref. [22]). (D) Average ^13^C and ^15^N chemical shift perturbation in Aβ_1-42_ fibrils with the presence of aducanumab, with the dotted line in the top panel corresponding to a threshold value of 0.25 ppm. Absence of bars reflect the lack of chemical shift assignment for those residues in either sample. (E) Structure of Aβ_1-42_ dimer (one protofilament stack) with the perturbed region highlighted in yellow. (F) Side-by-side representation of the N-terminal segment of Aβ_1-42_ fibrils without (top) and with (bottom) aducanumab, illustrating reduced dynamics upon antibody binding.

These results allow us to contrast the molecular modes of secondary nucleation by aducanumab and the BRICHOS domain from the lung surfactant protein C (Figure 6A-C). Secondary nucleation of Aβ_1-42_ seems to occur at small and rare sites ca. one site per 150 monomers in the fibril, likely corresponding to structural defects in the fibril core[25, 26]. These sites can be annealed by thermal treatment, reducing both nucleation efficiency and BRICHOS binding stoichiometry[26]. The current understanding of the complete elimination of secondary nucleation by BRICHOS[22] thus relies on binding to fibril defects with moderate affinity and low stoichiometry[25, 26]. This is confirmed by the minimal ^13^C and ^15^N chemical shift changes in ssNMR spectra of Aβ_M1-42_ fibrils formed in the presence of BRICHOS (Figure S9). More extensive, but still relatively small, chemical shift perturbations (including ^1^H shifts) were reported for Aβ_1-42_ fibrils formed in the presence of R221E mutant of Bri2-BRICHOS[56].

In contrast, aducanumab suppresses secondary nucleation only moderately (by a factor of ∼3 [27]) and likely acts through a different mechanism. Rather than binding to fibril defects, aducanumab appears to densely decorate the fibril surface, particularly along the exposed N-terminal domain. This may sterically hinder monomer access to catalytic sites or interfere with monomer conversion and detachment steps. Such physical hindrance of secondary nucleation may be amplified by the lateral association of fibrils, possibly governed by some antibody molecules (with its two binding sites) binding to N-termini on separate fibrils. This interpretation is supported by the observation of fibril bundling in the presence of aducanumab in cryo-TEM images (Figure 1 and S3). The observation of aducanumab binding to N-termini without interference with the fibril core (Figure 6D-F) further supports the view that the relatively high stoichiometry is limited only by the size of the antibody relative to the spacing on the N-termini of the fibrils[27]. In addition, although the intrinsic affinity for monomeric Aβ is low, aducanumab achieves high avidity through its two epitope-binding sites and the dense display of N-terminal epitopes along the fibril[39].

Simple geometric considerations further support this model: coating a fibril of ∼5 nm radius with an antibody layer (∼10 nm) increases the cross-sectional area (and volume for a given length) by nearly 9-fold, which is consistent with the observed 1:4.5 stoichiometry (∼7.2:1 mass ratio). This indicates that dense decoration of the fibril surface by aducanumab is physically plausible under our experimental conditions (Figure S7).

A third mode of action in the inhibition of secondary nucleation has been reported for α-synuclein and the R221E mutant of Bri2-BRICHOS, in which case inhibitor binding to the flexible C-termini that decorate α-synuclein fibrils hinders secondary nucleation, which in this system may occur at such fuzzy coat[57].

## Conclusion

Taken together, this work investigates the structural and dynamic changes of Aβ_1-42_ fibrils at atomic resolution upon antibody binding. We find that while the fibril core remains structurally unaltered, aducanumab engagement leads to significant perturbations in the dynamics of the N-terminal region. Additionally, perturbations are detected in the rigid C-terminal segment (G37–A42). These changes likely arise from its outward-facing position and spatial proximity to the antibody-bound N-termini of neighboring monomers, highlighting that antibody binding can influence both ends of the fibril without altering the core architecture. These changes are reflected in the disappearance of INEPT signals from flexible residues and the emergence of new or perturbed resonances in RFDR and DARR spectra, including signals from S8, Y10, histidine(s), and notably, D1 and E3. Furthermore, the experiments support a model in which aducanumab binds along the fibril surface and occasionally cross-links fibrils, as also observed by cryo-TEM bundling, thereby hindering for monomer to access catalytic defect sites on the fibril. However, because aducanumab does not directly target catalytic defect sites, secondary nucleation is only partially inhibited, in contrast to BRICHOS domains that completely attenuates secondary nucleation by binding to defects. Our results demonstrate the utility of high-resolution MAS NMR for resolving subtle yet functionally significant perturbations in aggregated protein assemblies. While our findings are specific to aducanumab, its binding mode, targeting the Aβ N-terminus with low intrinsic affinity, high avidity, and selectivity for aggregated forms of Aβ, is shared by several antibodies, including lecanemab and gantenerumab. These insights could help guide the design of improved antibody-based therapies for Alzheimer’s disease.

## Materials and Methods

### Sample Preparation

#### Generation of anti-Aβ antibody aducanumab

Aducanumab was generated as described in Sevigny, et al. [29] Briefly, the cDNAs for variable domains of aducanumab (V_H_ and V_L_) amino acid sequences were cloned into expression vectors containing hIgG1/kappa constant heavy (CH) chain and constant light (CL) chain domains separately. The two plasmids containing the full heavy chain or full light chain cDNAs were co-transfected into Chinese hamster ovary dihydrofolate reductase (*dhfr*) deficient host cell line, CHO-DG44I. The stably expressed antibody proteins were purified by protein-A affinity followed by ion-exchange chromatography.

#### Expression and purification of Aβ_1-42_ peptide

Aβ_1-42_ was expressed in fusion with the self-cleavable tag nPro [58] in the form of its EDDIE mutant [59]. The amino acid sequence, with the Aβ_1-42_ sequence underlined, is MELNHFELLYKTSKQKPVGVEEPVYDTAGRPLFGNPSEVHPQSTLKLPHDRGEDDIETTLRDLP RKGDCRSGNHLGPVSGIYIKPGPVYYQDYTGPVYHRAPLEFFDETQFEETTKRIGRVTGSDGKL YHIYVEVDGEILLKQAKRGTPRTLKWTRNTTNCPLWVTSC<u>DAEFRHDSGYEVHHQKLVFFAEDVGSNKGAIIGLMVGGVVIA</u>. The fusion protein was expressed from a Pet3a plasmid in M9 minimal medium prepared with ^13^C-glucose and ^15^NH_4_Cl as the sole carbon and nitrogen source, respectively. The purification involves the isolation of EDDIE-Aβ_1-42_ in urea-denatured form by anion exchange chromatography, autoproteolysis upon refolding of EDDIE to release Aβ_1-42_ and the purification of tag-free Aβ_1-42_ by anion exchange chromatography and size exclusion chromatography as described in detail [27]. The fractions dominated by Aβ_1-42_ monomer were lyophilized. Note that we have in previous work used a SUMO-fusion construct [60] for production of Aβ_1-42_; however, due to the need for expensive proteases and overlap elution of SUMO tag and Aβ_1-42_ in SEC, we have in this work used the self-cleavable EDDIE tag, which requires no protease and separates better from Aβ_1-42_. The cleaved off peptide is in both cases Aβ_1-42_ or native human sequence.

#### Sample preparation for MAS NMR

Lyophilized Aβ_1-42_ monomer samples from the same purification batch were dissolved in 10 mL, 6 M GuHCl and isolated from residual *E. coli* proteins, aggregates and small molecule contaminants using size exclusion chromatography on a Superdex 75 26/600 column (Figure S1). The eluted fractions were examined using UV absorbance, agarose gel electrophoresis and SDS PAGE with Coomassie staining. Fractions corresponding to the center of the Aβ_1-42_ monomer peak were collected in low-binding tubes and stored on ice until mixing with aducanumab and salt. Antibody aliquots from the same purification batch were further purified from remaining aggregates and lower MW contaminants using size exclusion chromatography on a 10×300 mm Superdex 200 column eluted in 20 mM NaP, 0.2 mM EDTA, pH 8.0, with 500 µl injected at a time (Figure S1). The peptide and antibody concentrations were determined by absorbance of the collected peaks using ε_280_ = 1490 M^-1^cm^-1^ and ε_280_ = 210000 M^-1^cm^-1^, respectively. The antibody was diluted 20 times before recording the spectrum to ensure that the absorbance was in the linear range. The Aβ_1-42_ monomer was supplemented with 100 mM NaCl from a 3 M NaCl stock in 20 mM NaP, 0.2 mM EDTA, pH 8.0, just prior to mixing with antibody in 4.5:1 molar ratio. Several such mixtures were prepared and pooled into the same glass bottle (Pyrex 500 mL, final concentrations 18 µM Aβ_1-42_ and 4 µM), which was incubated quiescent at room temperature for 4 days. An aliquot was withdrawn for cryo-TEM imaging to validate the formation of fibrils (Figure 1 and S3), before the sample was aliquoted in 50 mL falcon tubes and shipped at ambient temperature. The samples were then spun down into MAS rotors (3.2 mm for standard probe and 3.2 mm rotor for CryoProbe) using custom designed, home-built centrifuge tools in an ultracentrifuge at 30,000 RPM.

The Aβ_1-42_: Brichos samples were prepared using a similar procedure. Specifically, U-^13^C,^15^N-Aβ _M1-42_ was expressed and purified as described by Colvin, et al. [52, 61]. Unlabeled pro-SPC Brichos was expressed and purified as described by Willander, et al. [62].

### 800 MHz NMR Experiments

1D ^13^C and ^15^N cross-polarization (CP) (Figure 1) magic-angle-spinning (MAS) experiments were collected at 18.8 T (ω_1H_/2π=799.33 MHz ^1^H, ω_13C_/2π=200.99 MHz ^13^C) on a widebore Bruker Avance III spectrometer equipped with a 3.2 mm standard probe. Experiments were performed at a MAS rate of ω_r_/2π=20.0 kHz and at T=277 K. Cross polarization was accomplished with a ^13^C spin lock of 75.8 kHz and ^1^H spin lock of ω_1_/2π=59.5 kHz (linearly ramped ±15%) for 1 ms using ω_1_/2π=83.0 kHz TPPM20 ^1^H decoupling was used during detection. 2D ^13^C-^13^C RFDR correlation experiment (Figure S6A) were performed at a MAS rate of ω_r_/2π=20.0 kHz. Cross Polarization was accomplished with a ^13^C spin lock of ω_1_/2π=75.8 kHz and ^1^H spin lock of ω_1_/2π=59.5 kHz (linearly ramped ±15%). The RFDR mixing time was 1.6 ms and the π-pulses used ω_1_/2π=75.8 kHz RF amplitude. The decoupling during acquisition and t1 evolution were TPPM20 using ω_1_/2π=83.0 kHz ^1^H RF amplitudes, while the decoupling during RFDR mixing was ω_1_/2π=71.4 kHz CW decoupling. In the indirect dimension, 512 complex points with a dwell of 25.0 μs (spectral width ∼200 ppm) were acquired. A recycling delay of 4 s was used for all the above experiments. These parameters were typically used for acquisition of the Aβ_1-42_, Aβ_1-42_:Adu, and Aβ_1-42_:Brichos spectra.

### CPMAS CryoProbe 600 MHz NMR Experiments

1D ^13^C and ^15^N cross-polarization (CP) magic-angle-spinning (MAS) experiments (Figure 4) were collected at 14.1 T (ω_1H_/2π=600.16 MHz ^1^H, ω_13C_/2π=150.91 MHz ^13^C) on a standard bore Bruker NEO spectrometer equipped with a 3.2 mm CPMAS cryoprobe (Bruker Biosolids CryoProbe™) in which the detection coil and preamplifier operate at cryogenic temperatures to increase sensitivity while the sample remains at lab temperature. Experiments were performed at a MAS rate of ω_r_/2π=13.5 kHz. Cross polarization was accomplished with a ^13^C spin lock of ω_1_/2π=55 kHz and ^1^H spin lock ω_1_/2π=68 kHz (linearly ramped ±5%) and a ^15^N spin lock of ω_1_/2π =37 kHz and ^1^H spin lock of ω_1_/2π=50.5 kHz (linearly ramped ±5%), respectively. Spinal64 ^1^H decoupling [63] with ω_1_/2π= 83 kHz was used during detection. ^13^C spectra consist of the sum of 4,096 transients acquired with a relaxation delay of 4 s, for a total acquisition time of 4.5 h. ^13^C chemical shifts were referenced indirectly to DSS via an external solid-state sample of adamantane with the downfield-shifted peak set to 40.48 ppm [64, 65]. ^15^N spectra consist of the sum of 16,384 transients acquired with a relaxation delay of 4 s, for a total acquisition time of 18 h. ^15^N chemical shifts were referenced indirectly to liq-NH_3_ (25 °C) via the frequency ratio method [65]. The sample temperature was regulated with a Bruker variable temperature controller set to 5 °C.

2D ^13^C-^13^C RFDR [66, 67] and DARR [68] correlation experiments (Figures 2 and 3) were collected at 14.1 T ((ω_1H_/2π=600.16 MHz ^1^H, ω_13C_/2π=150.91 MHz ^13^C) on a standard bore Bruker NEO spectrometer equipped with a 3.2 mm CPMAS cryoprobe (Bruker Biosolids CryoProbe™). Experiments were performed at a MAS rate of 13.5 kHz. Cross Polarization was accomplished with a ^13^C spin lock of ω_1_/2π=55 kHz and ^1^H spin lock of ω_1_/2π=68 kHz (linearly ramped ±5%). ^13^C and ^1^H pulses of ω_1_/2π=50 kHz and 83 kHz were used throughout and ω_1_/2π=83 kHz Spinal64 ^1^H decoupling [63] was used during detection. The RFDR mixing time was 1.6 ms and the DARR mixing time was 100 ms. In the indirect dimension, 512 complex points with a dwell of 27.4 μs (spectral width 40.486 kHz, total acquisition time 12.64 ms) were acquired. In the direct dimension, 1024 complex points with a dwell of 20 μs (spectral width 50 kHz, total acquisition time 20.48 ms) were acquired. For the RFDR experiments, 48 transients at each t_1_ point were coadded with a relaxation delay 4 s for a total experiment time of 58 hr. For the DARR experiments, 80 transients at each t_1_ point were coadded with a relaxation delay 4 s for a total experiment time of 97 hr. The sample temperature was regulated with a Bruker variable temperature controller set to 5 °C.

The pulse sequences used are represented in Figure S4.

### Cryo-TEM Imaging

Samples for cryo-TEM were taken after 4 days incubation. Specimens for cryo-TEM were prepared in an automatic plunge freezer system (Leica EM GP). The climate chamber temperature was kept at 21°C, and relative humidity was ≥90% to minimise loss of solution during sample preparation. 4 µL samples were loaded onto 300 mesh lacey carbon-filmed copper TEM grids and blotted with a filter paper to absorb extra solution. The grid was then plunged into liquid ethane (-180 °C) to flash freeze all samples and stored in liquid nitrogen until imaged on the next day. A Fischione Model 2550 cryo transfer tomography holder was used to transfer the specimen into the electron microscope, JEM 2200FS, equipped with an in-column energy filter (Omega filter), which allows zero-loss imaging. The acceleration voltage was 200kV and zero-loss images were recorded digitally with a TVIPS F416 camera using SerialEM under low dose conditions with a 10 eV energy selecting slit in place.

## Acknowledgments

The authors thank Neurimmune for providing access to aducanumab for this study. This research was supported by grants from the National Institute of Aging (AG058504) to RGG, the National Institute of General Medical Sciences to LJM (GM145369) and to RS (R35GM142912), and by the Swedish Research Council, VR2015-00143 to SL. FQ and PW were supported by Biogen, Inc. The 600 MHz NMR spectrometer and MAS Cryoprobe were purchased with funds from NIH grant S10OD032197 to LJM.

