## Supporting Information for "Aducanumab Binding to Aβ1-42 Fibrils Alters Dynamics of the N-Terminal Tail While Preserving the Fibril Core"

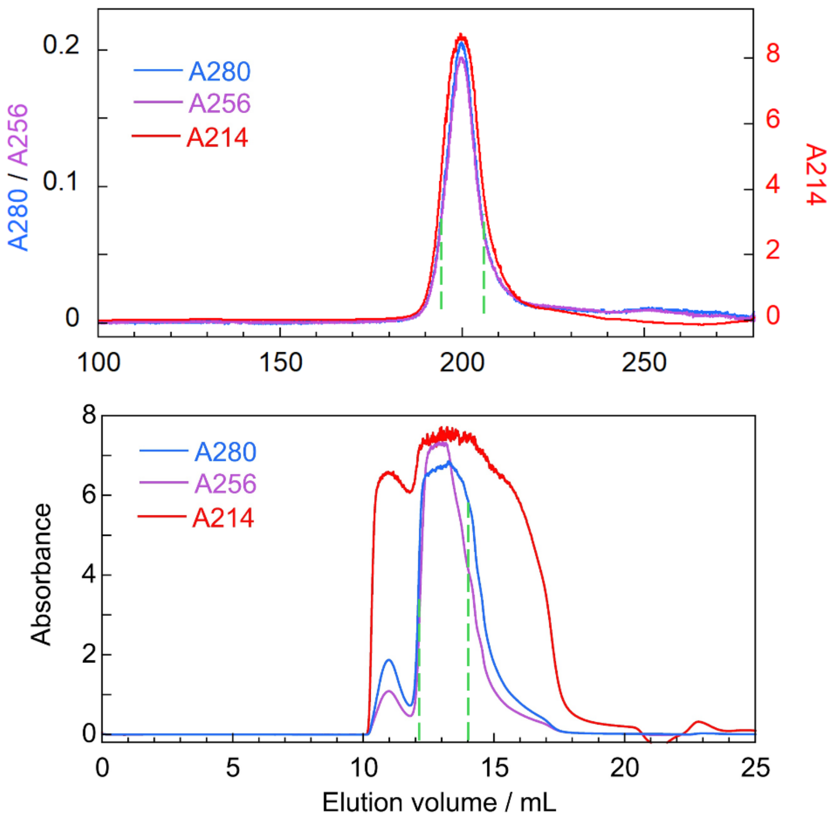

**Figure S1.** Examples of size exclusion chromatograms from the final isolation of A$\beta_{1\text{-}42}$ (top) and aducanumab (bottom) before mixing to prepare samples for MAS NMR shown as the absorbance with 10 mm path length. Fractions between green dashed lines were collected. The aducanumab signals reach far above the threshold of the detector on the chromatography system and the absorbance spectrum used for concentration determination was therefore measured in a 10 mm path-length quartz cuvette using a Labbot instrument (Probation Labs Sweden AB) after 20-fold dilution of a small part of the collected fraction.

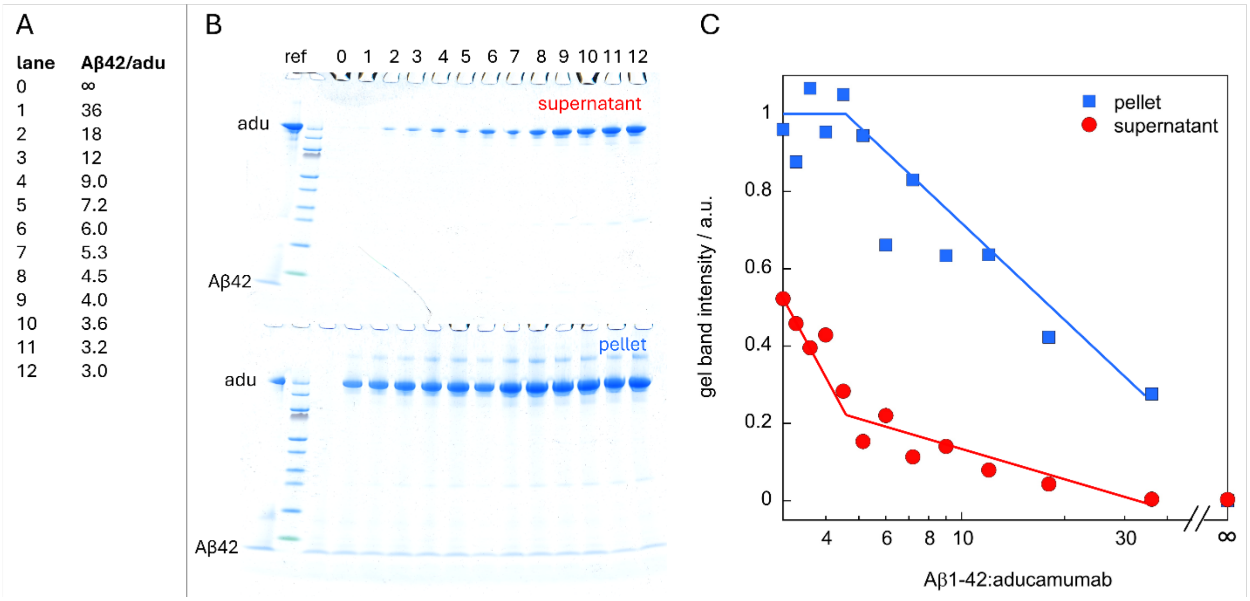

**Figure S2**. Sedimentation assay. Samples of $A\beta_{1\text{-}42}$ at constant concentration 18 μM and aducanumab at varying concentration 0 or 0.5-6 μM in steps of 0.5 μM in 20 mM NaP, 0.2 mM EDTA, 100 mM NaCl, pH 8.0, were incubated for 3 days at room temperature in low-binding tubes (Axygen) at quiescent conditions followed by centrifugation at 20,000 g for 10 min. The pellet and supernatant from each tube were mixed 1:1 with using non-reducing loading buffer and analyzed by SDS PAGE on a 10-20% Tris/Tricine gel (Novex) alongside a non-aggregated reference sample. (A) molar ratio of $A\beta_{1\text{-}42}$ and aducanumab in the sample loaded in each lane 0-12. (B) Stained gels (Coomassie quick stain) with the migration of aducanumab (adu) and $A\beta_{1\text{-}42}$ indicated to the left. (C) The gel band intensity of aducanumab from each lane, estimated using ImageJ, *versus* molar ratio $A\beta_{1\text{-}42}$:aducanumab. The solid lines indicate the break point in behavior close to a molar ratio of 4.5:1.

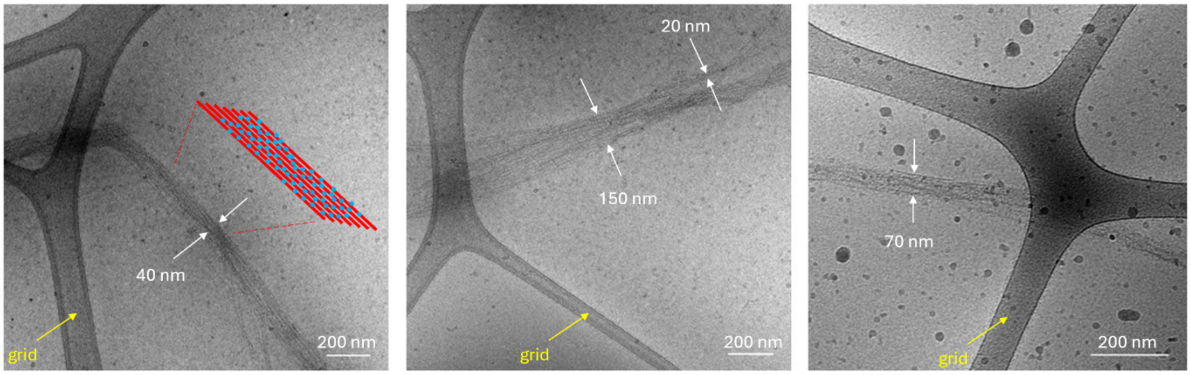

**Figure S3**. Additional cryo-EM images of fibrils formed from A$\beta_{1-42}$ in the presence of aducanumab collected at low magnifications showing wider fields compared to the images in the main text. The scale bar in each panel is 200 nm.

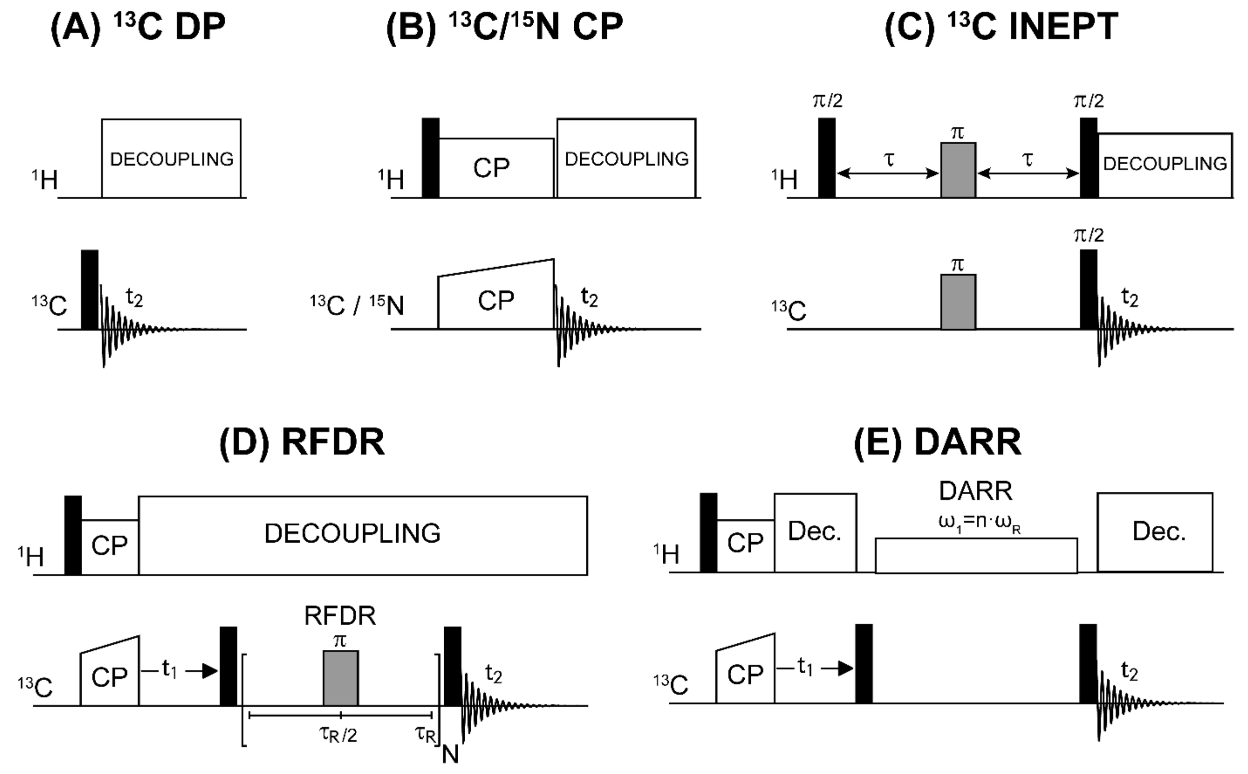

**Figure S4.** Pulse sequences for the MAS NMR experiments used in this study. (A) $^{13}C$ Direct Polarization (DP) detects signals from both rigid and mobile residues. (B) $^{13}C$ Cross Polarization (CP) experiment detects signals from rigid residues via dipolar coupling. (C) $^{13}C$ INEPT (Insensitive Nuclear Enhancement by Polarization Transfer) detects signals from mobile residues via scalar (J) coupling, sensitive to motions on the nanosecond timescale or faster. (D) RFDR (Radiofrequency-Driven Dipolar Recoupling) experiment probes short-range $^{13}C$–$^{13}C$ contacts among rigid residues. (E) DARR (Dipolar Assisted Rotational Resonance) probes long-range $^{13}C$–$^{13}C$ contacts among rigid residues using continuous-wave $^{1}H$ irradiation.

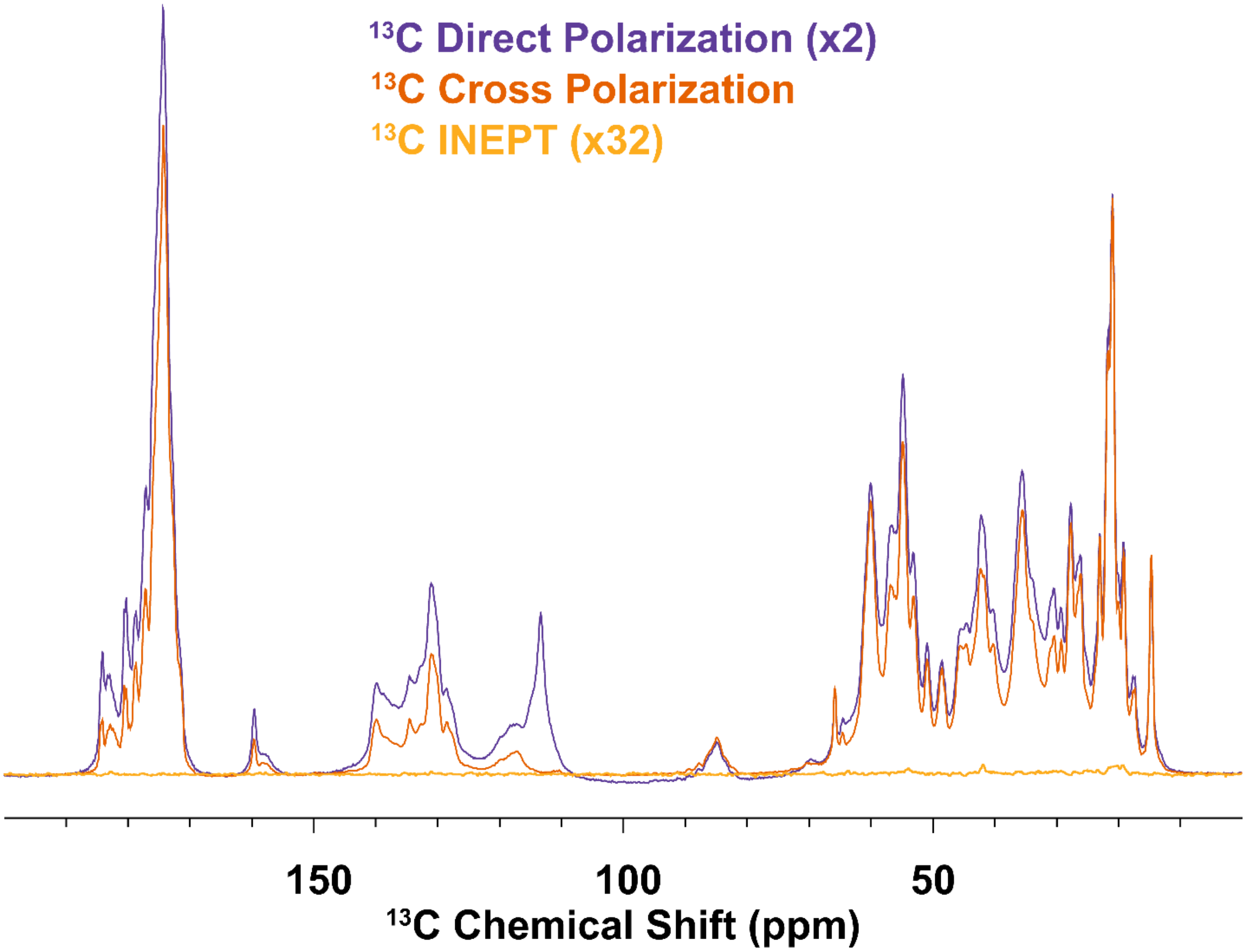

**Figure S5.** Comparison of $^{13}C$ NMR signal intensities of the $A\beta_{1-42}$ sample prepared in the presence of aducanumab acquired using three different polarization transfer methods: direct $^{13}C$ polarization (DP, purple, scaled x2), $^{1}H$–$^{13}C$ cross-polarization (CP, orange), and $^{1}H$–$^{13}C$ INEPT (yellow, scaled x32). The DP spectrum is scaled by a factor of 2 to account for the difference in gyromagnetic ratio (x4) and the typical CP efficiency (~0.5), i.e., 4 x 0.5 = 2. INEPT shows only weak signals (scaled x32 to highlight the lack of intensity), indicating that few residues retain sufficient mobility for scalar-coupling (J) based transfer. The aliphatic and carbonyl regions show comparable intensities between DP and CP, whereas the aromatic side chains appear stronger in DP than in CP but are absent in INEPT. This suggests that these side chains undergo intermediate timescale motions: too rigid to be detected by INEPT, yet too dynamic to be fully captured by CP.

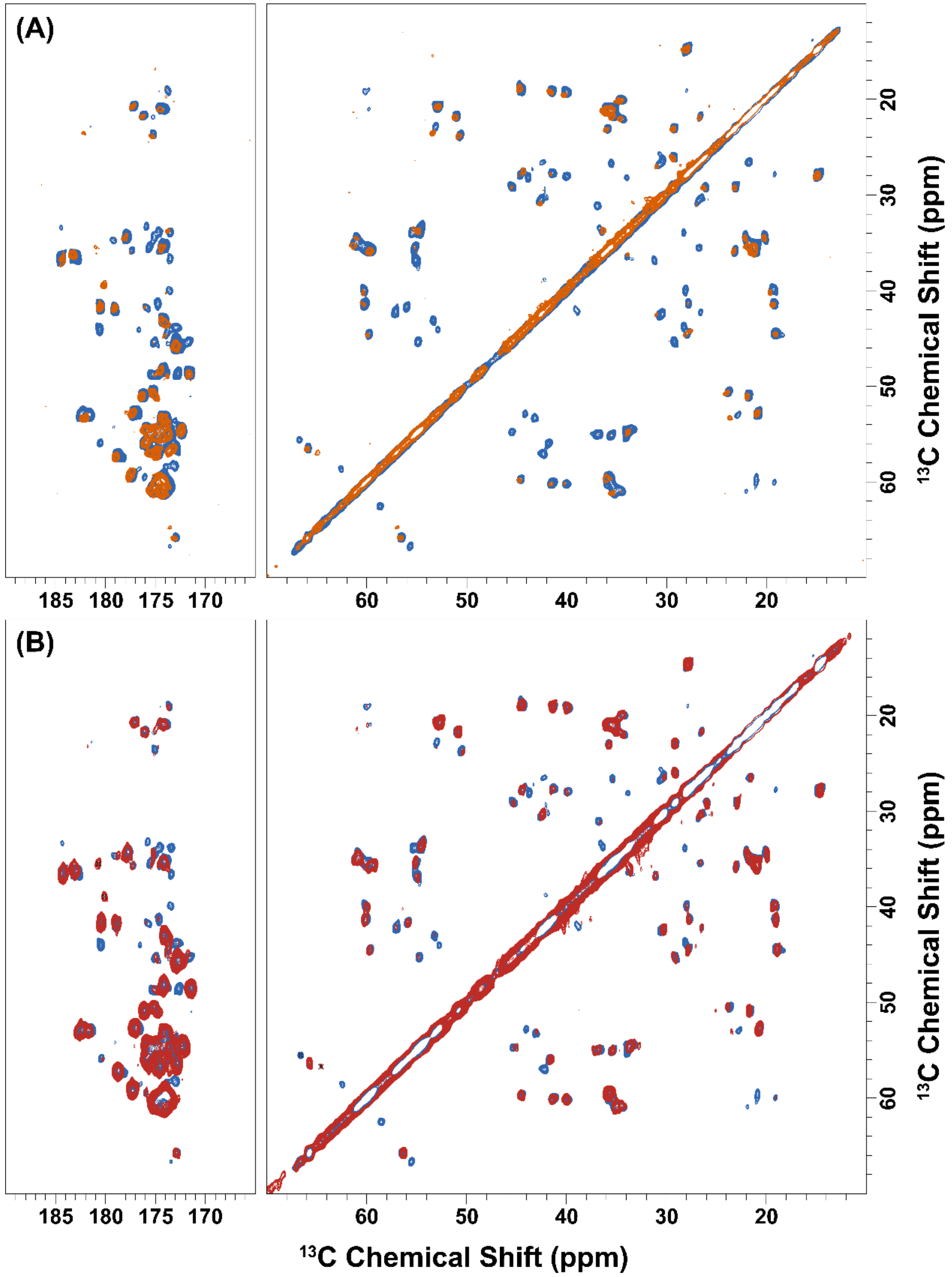

**Figure S6.** Comparison of signal intensities in 2D $^{13}C$–$^{13}C$ RFDR spectra of Aβ$_{1-42}$ fibrils in the presence of aducanumab, acquired using different probes. (A) Spectrum recorded at 800 MHz using a standard 3.2 mm MAS probe (orange). (B) Spectrum recorded at 600 MHz using a 3.2 mm MAS CryoProbe (red). In both panels, the overlaid blue spectra correspond to unbound Aβ$_{1-42}$ fibrils from Silvers et al [1]. The CryoProbe spectrum (B) shows enhanced sensitivity, enabling the detection of weaker peaks not observed in the standard probe spectrum (A).

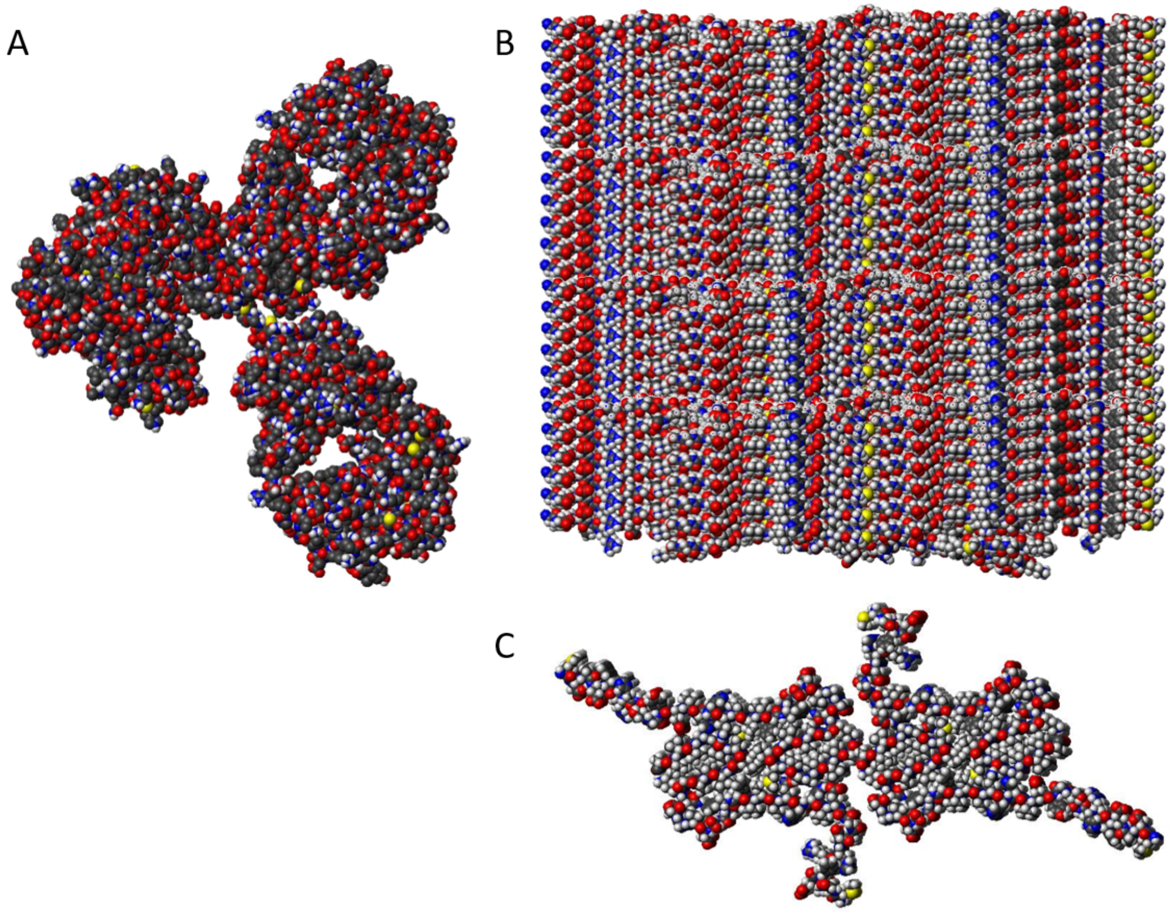

**Figure S7**. Size comparison of an IgG antibody (A, 1igy.pdb) and A$\beta_{1\text{-}42}$ fibril (B,C) displaying 24 planes of the A$\beta_{M01\text{-}42}$ fibril structure derived from SAXS and ssNMR data[2] with four monomers per fibril plane viewed from the side (B) and top (C). The image was prepared as a CPK model using Molmol[3].

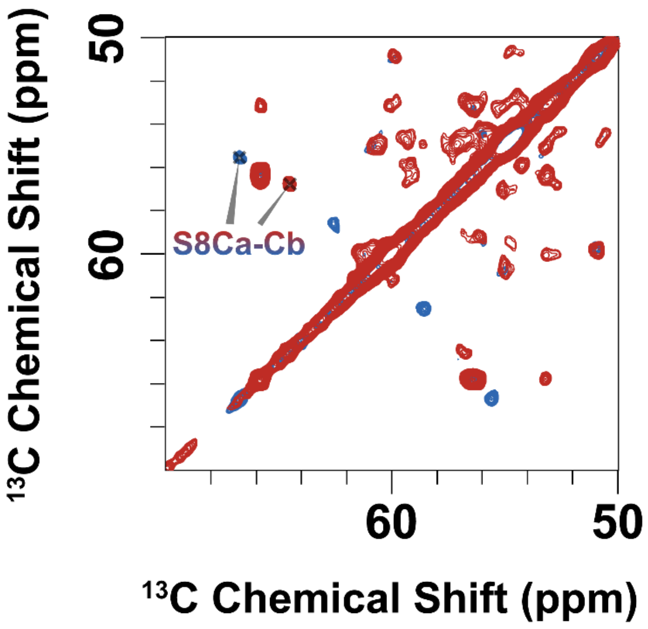

**Figure S8.** Expanded view of S8 perturbation in 100 ms DARR (Figure 4 in the main text).

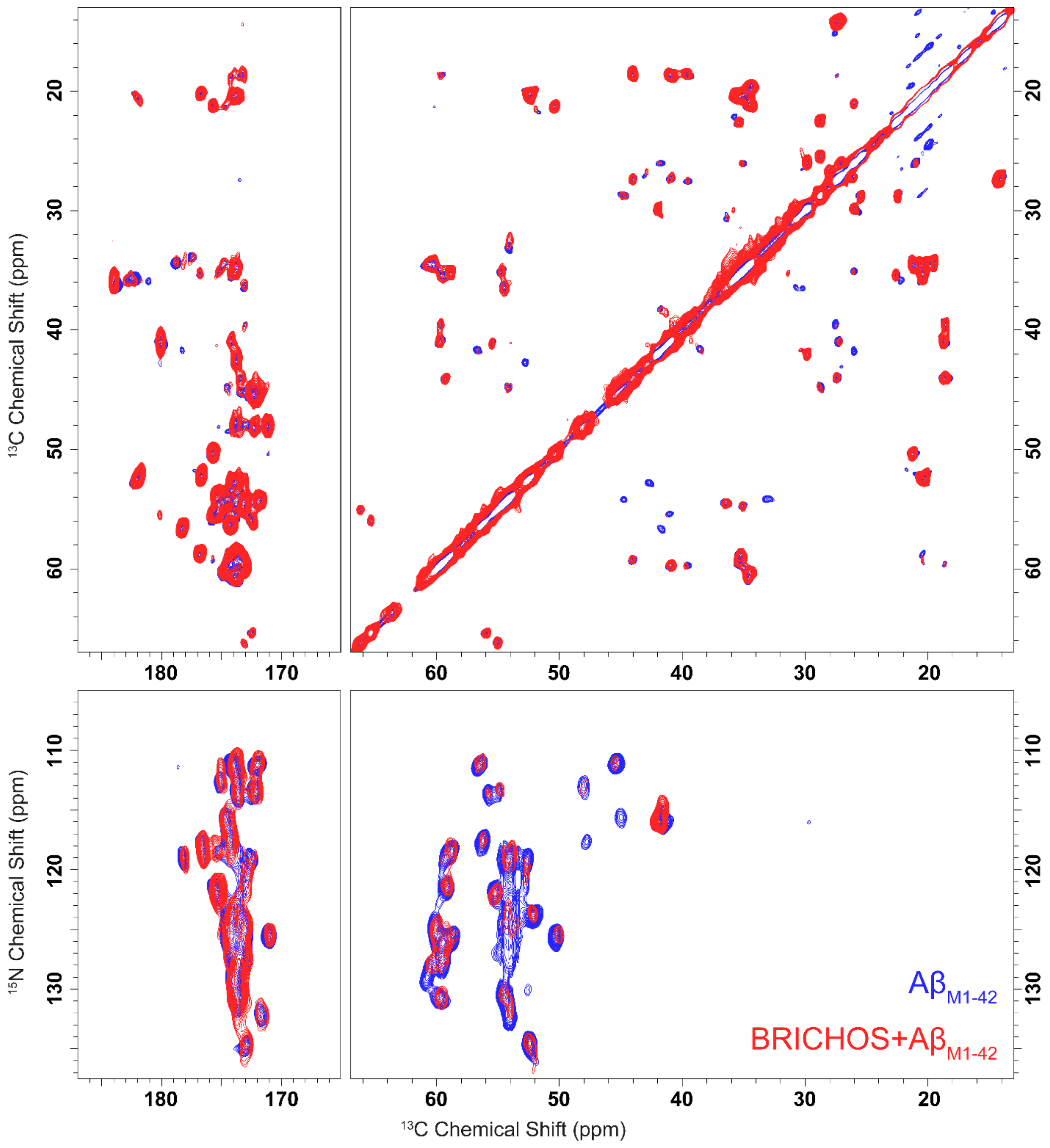

**Figure S9.** 1.6 ms mixing $^{13}C$-$^{13}C$ RFDR spectrum (top panels) and 1.6 ms mixing $^{15}N$-$^{13}C$ ZF-TEDOR (bottom panels) recorded at 800 MHz, $\omega_r$ =20 kHz on Aβ$_{M1-42}$ (blue) and Aβ$_{M1-42}$ in the presence of BRICHOS.

**Table S1.** $^{13}C$ and $^{15}N$ chemical shift assignments of $A\beta_{1\text{-}42}$ fibrils reproduced from ref [4]. Note: Chemical shifts of A2, S8b, G9, G38 have been updated from the previous reported values upon examining the data. A second set of cross peaks corresponding to Ser8 (structural heterogeneity) was also observed in the original dataset, which disappears upon aducanumab binding.

| Residue | Res Type | N | C | CA | CB | CG | CG1 | CG2 | CD | CD1 | CD2 | CE | CZ | N |
|---|---|---|---|---|---|---|---|---|---|---|---|---|---|---|
| 1 | D | | | 54.28 | 42.26 | | | | | | | | | |
| 2 | A | | | 50.68 | 23.84 | | | | | | | | | |
| 3 | E | | | 56.49 | 29.83 | 35.91 | | | 183.3 | | | | | |
| 4 | F | | | 55.02 | 39.11 | 120 | | | 131.4 | | | 137.6 | 129.7 | |
| 5 | R | 121.9 | 176.5 | 54.09 | 33.14 | 27.1 | | | | | | | 159.22 | |
| 6 | H | | | | | | | | | | | | | |
| 7 | D | | 172.16 | | | | | | | | | | | |
| 8 | S | 113.2 | 172.94 | 55.07 | 66.14 | | | | | | | | | |
| 8b | S | 114.24 | 176.91 | 58.74 | 62.68 | | | | | | | | | |
| 9 | G | 108.69 | 174.18 | 44.00 | | | | | | | | | | |
| 10 | Y | 113 | 176.8 | 55 | 41.35 | 135.8 | | | 132.4 | | | 117.2 | 156.4 | |
| 11 | E | 120.5 | 173.7 | 52.54 | 35.66 | 32.43 | | | 182.1 | | | | | |
| 12 | V | 124.6 | 172.1 | 60.25 | | | | | | | | | | |
| 13 | H | | | | | | | | | | | | | |
| 14 | H | | | | | | | | | | | | | |
| 15 | Q | | | | | | | | | | | | | |
| 16 | K | 124.43 | 171.78 | 54.53 | | 25.71 | | | 30.18 | | | | | |
| 17 | L | 132.09 | 174.41 | 54.2 | 44.76 | 28.71 | | | | 22.44 | 25.45 | | | |
| 18 | V | 125.23 | 173.71 | 58.94 | 35.19 | 20.30 | | | | | | | | |
| 19 | F | 130.35 | 174.38 | 59.89 | 40.29 | 139.61 | | | 130.50 | | | 129.67 | | |
| 20 | F | 117.18 | 174.24 | 56.33 | 40.25 | 139.37 | | | 130.60 | 129.93 | | | | |
| 21 | A | 123.47 | 176.69 | 52.25 | 20.28 | | | | | | | | | |
| 22 | E | 118.54 | 175.34 | 54.03 | 33.06 | 35.85 | | | 183.60 | | | | | |
| 23 | D | 121.95 | 175.36 | 55.42 | 40.98 | 179.95 | | | | | | | | |
| 24 | V | 117.95 | 176.78 | 58.86 | 35.32 | | 20.40 | 22.56 | | | | | | |
| 25 | G | 117.21 | 173.73 | 47.71 | | | | | | | | | | |
| 26 | S | 113.29 | 172.41 | 55.93 | 65.34 | | | | | | | | | |
| 27 | N | 118.53 | 173.76 | 52.81 | 42.70 | | | | | | | | | |
| 28 | K | 130.25 | 175.17 | 54.76 | 35.04 | 26.00 | | | 29.81 | | | 42.04 | | 34.00 |
| 29 | G | 112.48 | 171.06 | 48.16 | | | | | | | | | | |
| 30 | A | 125.09 | 175.72 | 50.25 | 21.26 | | | | | | | | | |
| 31 | I | 121.06 | 173.38 | 59.30 | 44.02 | | 27.35 | 18.65 | | 14.07 | | | | |
| 32 | I | 125.74 | 174.12 | 59.67 | 40.95 | | 27.32 | 18.64 | | 14.11 | | | | |
| 33 | G | 110.76 | 172.22 | 45.39 | | | | | | | | | | |
| 34 | L | 110.68 | 178.26 | 56.70 | 41.70 | 27.14 | | | | 20.92 | 25.99 | | | |
| 35 | M | 118.64 | 172.99 | 54.48 | 36.38 | 30.62 | | | | | | 17.00 | | |
| 36 | V | 124.60 | 174.68 | 60.15 | 34.52 | | 19.49 | 21.22 | | | | | | |
| 37 | G | 115.37 | 173.30 | 45.03 | | | | | | | | | | |
| 38 | G | 115.20 | 173.10 | 49.22 | | | | | | | | | | |
| 39 | V | 129.15 | 174.50 | 61.13 | 34.67 | | 20.22 | 21.13 | | | | | | |
| 40 | V | 128.18 | 173.77 | 60.83 | 34.56 | | 20.70 | 20.32 | | | | | | |
| 41 | I | 127.45 | 173.03 | 59.58 | 39.48 | | 27.40 | 18.61 | | 14.18 | | | | |

| | | | | | |
|---|---|---|---|---|---|
| 42a | A | 134.14 | 181.60 | 53.06 | 22.86 |
| 42b | A | 134.14 | 182.08 | 52.51 | 20.28 |

**Table S2.** $^{13}C$ and $^{15}N$ chemical shift assignments of Aβ$_{1\text{-}42}$ fibrils prepared in the presence of aducanumab.

| Residue | Res Type | N | C | CA | CB | CG | CG1 | CG2 | CD | CD1 | CD2 | CE | CZ | N |
|---|---|---|---|---|---|---|---|---|---|---|---|---|---|---|
| 1 | D | | 171.68 | 53.82 | 41.18 | 178.53 | | | | | | | | |
| 2 | A | 119.02 | 174.75 | 50.08 | 23.40 | | | | | | | | | |
| 3 | E | 119.65 | | 54.31 | 29.81 | 35.69 | | | 182.73 | | | | | |
| 4 | F | | | | | | | | | | | | | |
| 5 | R | | | 54.67 | 33.10 | 26.65 | | | 43.65 | | | | 159.23 | |
| 6 | H | | | | | | | | | | | | | |
| 7 | D | | 173.59 | 53.61 | 42.60 | | | | | | | | | |
| 8 | S | 111.39 | 173.09 | 56.18 | 64.04 | | | | | | | | | |
| 9 | G | 114.06 | 173.11 | 42.79 | | | | | | | | | | |
| 10 | Y | | | | | 123.15 | | | 132.09 | | | 117.75 | 156.97 | |
| 11 | E | | | | | | | | | | | | | |
| 12 | V | 124.70 | 173.36 | 60.26 | 34.68 | | 20.98 | 19.52 | | | | | | |
| 13 | H | | | | | | | | | | | | | |
| 14 | H | | | | | | | | | | | | | |
| 15 | Q | | | | 36.15 | 33.98 | | | 177.41 | | | | | |
| 16 | K | | 171.83 | 54.23 | | 25.98 | | | | | | 41.19 | | |
| 17 | L | 131.24 | 174.42 | 54.28 | 44.89 | 28.70 | | | | 22.44 | 25.56 | | | |
| 18 | V | 125.44 | 173.71 | 59.13 | 35.08 | 20.48 | | | | | | | | |
| 19 | F | 130.29 | 174.25 | 59.67 | 40.57 | 139.66 | | | 130.23 | | | 129.51 | | |
| 20 | F | 117.17 | 174.08 | 56.34 | 40.01 | 139.49 | | | 130.47 | | | 129.14 | 126.75 | |
| 21 | A | 123.56 | 176.63 | 52.25 | 20.30 | | | | | | | | | |
| 22 | E | 118.52 | 175.37 | 54.15 | 33.14 | 35.83 | | | 183.82 | | | | | |
| 23 | D | 121.79 | 175.34 | 55.37 | 41.19 | 179.99 | | | | | | | | |
| 24 | V | 118.16 | 176.84 | 58.93 | 35.27 | | 20.42 | 22.51 | | | | | | |
| 25 | G | 117.27 | 173.72 | 47.71 | | | | | | | | | | |
| 26 | S | 113.63 | 172.44 | 55.87 | 65.33 | | | | | | | | | |
| 27 | N | 118.85 | 173.62 | 52.71 | 42.61 | | | | | | | | | |
| 28 | K | 130.68 | 174.82 | 54.49 | 34.95 | 26.09 | | | 29.90 | | | 42.02 | | 34.00 |
| 29 | G | 112.66 | 171.02 | 48.08 | | | | | | | | | | |
| 30 | A | 125.28 | 175.66 | 50.43 | 21.23 | | | | | | | | | |
| 31 | I | 121.16 | 173.37 | 59.38 | 44.10 | | 27.38 | 18.58 | | 14.16 | | | | |
| 32 | I | 125.84 | 173.87 | 59.59 | 40.97 | | 27.35 | 18.62 | | 14.16 | | | | |
| 33 | G | 110.86 | 172.31 | 45.31 | | | | | | | | | | |
| 34 | L | 111.15 | 178.36 | 56.76 | 41.80 | 26.99 | | | | 21.17 | 26.08 | | | |
| 35 | M | 118.15 | 172.95 | 54.41 | 36.45 | 30.68 | | | | | | 16.94 | | |
| 36 | V | 124.56 | 174.59 | 60.24 | 34.58 | | 19.59 | 21.27 | | | | | | |
| 37 | G | 115.89 | 172.33 | 45.08 | | | | | | | | | | |
| 38 | G | 114.60 | 172.64 | 48.27 | | | | | | | | | | |
| 39 | V | | 173.72 | 60.42 | 34.84 | 20.59 | | | | | | | | |
| 40 | V | 127.37 | 173.74 | 60.39 | 34.92 | 20.58 | | | | | | | | |

| | | | | | | | | | | |
|---|---|---|---|---|---|---|---|---|---|---|
| 41a | I | 127.81 | 173.33 | 59.58 | 39.61 | | 27.45 | 18.70 | | 14.22 |
| 41b | I | 127.81 | 172.94 | 59.58 | 39.61 | | 27.45 | 18.70 | | 14.22 |
| 42a | A | 132.79 | 181.77 | 52.68 | 23.11 | | | | | |
| 42b | A | 134.24 | 182.24 | 52.59 | 20.39 | | | | | |

**Table S3.** Chemical shift differences between Aβ$_{1\text{-}42}$ fibrils prepared in the presence and absence of aducanumab. Values are color-coded to emphasize the magnitude of the changes.

| Residue | Res Type | N | C | CA | CB | CG | CG1 | CG2 | CD | CD1 | CD2 | CE | CZ | N |
|---|---|---|---|---|---|---|---|---|---|---|---|---|---|---|
| 1 | D | | | -0.46 | -1.08 | | | | | | | | | |
| 2 | A | | | -0.6 | -0.44 | | | | | | | | | |
| 3 | E | | | -2.18 | -0.02 | -0.22 | | | -0.57 | | | | | |
| 4 | F | | | | | | | | | | | | | |
| 5 | R | | | 0.58 | -0.04 | -0.45 | | | | | | | 0.01 | |
| 6 | H | | | | | | | | | | | | | |
| 7 | D | | 1.43 | | | | | | | | | | | |
| 8 | S | -1.81 | 0.15 | 1.11 | -2.1 | | | | | | | | | |
| 9 | G | 5.37 | -1.07 | -1.22 | | | | | | | | | | |
| 10 | Y | | | | | | | | -0.31 | | | 0.55 | 0.57 | |
| 11 | E | | | | | | | | | | | | | |
| 12 | V | 0.1 | 1.26 | 0.01 | | | | | | | | | | |
| 13 | H | | | | | | | | | | | | | |
| 14 | H | | | | | | | | | | | | | |
| 15 | Q | | | | | | | | | | | | | |
| 16 | K | | 0.05 | -0.3 | | 0.27 | | | | | | | | |
| 17 | L | -0.85 | 0.01 | 0.08 | 0.13 | -0.01 | | | | 0 | 0.11 | | | |
| 18 | V | 0.21 | 0 | 0.19 | -0.11 | 0.18 | | | | | | | | |
| 19 | F | -0.06 | -0.13 | -0.22 | 0.28 | 0.05 | | | -0.27 | | | -0.16 | | |
| 20 | F | -0.01 | -0.16 | 0.01 | -0.24 | 0.12 | | | -0.13 | | | | | |
| 21 | A | 0.09 | -0.06 | 0 | 0.02 | | | | | | | | | |
| 22 | E | -0.02 | 0.03 | 0.12 | 0.08 | -0.02 | | | 0.22 | | | | | |
| 23 | D | -0.16 | -0.02 | -0.05 | 0.21 | 0.04 | | | | | | | | |
| 24 | V | 0.21 | 0.06 | 0.07 | -0.05 | | 0.02 | -0.05 | | | | | | |
| 25 | G | 0.06 | -0.01 | 0 | | | | | | | | | | |
| 26 | S | 0.34 | 0.03 | -0.06 | -0.01 | | | | | | | | | |
| 27 | N | 0.32 | -0.14 | -0.1 | -0.09 | | | | | | | | | |
| 28 | K | 0.43 | -0.35 | -0.27 | -0.09 | 0.09 | | | 0.09 | | | -0.02 | | |
| 29 | G | 0.18 | -0.04 | -0.08 | | | | | | | | | | |
| 30 | A | 0.19 | -0.06 | 0.18 | -0.03 | | | | | | | | | |
| 31 | I | 0.1 | -0.01 | 0.08 | 0.08 | | 0.03 | -0.07 | | 0.09 | | | | |
| 32 | I | 0.1 | -0.25 | -0.08 | 0.02 | | 0.03 | -0.02 | | 0.05 | | | | |
| 33 | G | 0.1 | 0.09 | -0.08 | | | | | | | | | | |
| 34 | L | 0.47 | 0.1 | 0.06 | 0.1 | -0.15 | | | | 0.25 | 0.09 | | | |
| 35 | M | -0.49 | -0.04 | -0.07 | 0.07 | 0.06 | | | | | | -0.06 | | |
| 36 | V | -0.04 | -0.09 | 0.09 | 0.06 | | 0.1 | 0.05 | | | | | | |
| 37 | G | 0.52 | -0.97 | 0.05 | | | | | | | | | | |

| | | | | | | | | | | |
|---|---|---|---|---|---|---|---|---|---|---|
| 38 | G | -0.6 | -0.46 | -0.95 | | | | | | |
| 39 | V | | -0.78 | -0.71 | 0.17 | | | | | |
| 40 | V | -0.81 | -0.03 | -0.44 | 0.36 | | | | | |
| 41a | I | 0.36 | 0.3 | 0 | 0.13 | | 0.05 | 0.09 | | 0.04 |
| 41b | I | 0.36 | -0.09 | 0 | 0.13 | | 0.05 | 0.09 | | 0.04 |
| 42a | A | -1.35 | 0.17 | -0.38 | 0.25 | | | | | |
| 42b | A | 0.1 | 0.16 | 0.08 | 0.11 | | | | | |

| Experiment | MAS (kHz) | Transfer | $^{1}H$ Spin Lock (kHz) | $^{13}C$ Spin Lock (kHz) | $^{15}N$ Spin Lock (kHz) | Mixing Time (ms) | Decoupling (kHz / scheme) | Complex Points (t1, indirect dim) | Total Time (h) |
|---|---|---|---|---|---|---|---|---|---|
| 1D $^{13}C$ CP (799.33 MHz) | 20.0 | $^{1}H$–$^{13}C$ CP | 59.5 ±15% | 75.8 | – | 1.0 | 83 (TPPM20) | – | ~4.0 |
| 1D $^{15}N$ CP (799.33 MHz) | 20.0 | $^{1}H$–$^{15}N$ CP | 59.5 ±15% | – | 29.9 | 1.5 | 83 (TPPM20) | – | 16.0 |
| 2D $^{13}C$–$^{13}C$ RFDR (799.33 MHz) | 20.0 | CP–RFDR | 59.5 ±15% (CP) | 75.8 (CP) | – | 1.6 (RFDR; 75.8 kHz pi pulse) | 71.4 (CW during RFDR); 83 (TPPM20 during t1 and acquisition) | 512 (x25 us) | ~53 |
| 1D $^{13}C$ CP (600.16 MHz) | 13.5 | $^{1}H$–$^{13}C$ CP | 68 ±5% | 55 | – | – | 83 (SPINAL64) | – | 4.5 |
| 1D $^{15}N$ CP (600.16 MHz) | 13.5 | $^{1}H$–$^{15}N$ CP | 50.5 ±5% | – | 37 | – | 83 (SPINAL64) | – | 18 |
| 2D $^{13}C$–$^{13}C$ RFDR (600.16 MHz) | 13.5 | CP–RFDR | 68 (±5%) | 55 | – | 1.6 (RFDR; 50 kHz pi pulse) | 83 (SPINAL64) | 512 (x27.4 us) | 58 |
| 2D $^{13}C$–$^{13}C$ DARR (600.16 MHz) | 13.5 | CP–DARR | 68 (±5%) | 55 | – | 100 (DARR) | 83 (SPINAL64) | 512 (x27.4 us) | 97 |

**Table S4.** Experimental parameters for spectra shown in the main text.
